# Contribution of 6mer seed toxicity to HIV-1 induced cytopathicity

**DOI:** 10.1101/2022.10.01.510471

**Authors:** Aparajitha Vaidyanathan, Harry E. Taylor, Thomas J. Hope, Richard T. D’Aquila, Elizabeth T. Bartom, Judd F. Hultquist, Marcus E. Peter

## Abstract

HIV-1 (HIV) infects CD4 positive T cells, the gradual depletion of which can lead to the onset of Acquired Immunodeficiency Syndrome (AIDS) in the absence of antiretroviral therapy (ART). Several forms of cell death have been shown to be involved in HIV-mediated killing of either directly infected or bystander cells. It is still unknown, however, why some cells survive HIV infection and persist as part of the latently infected reservoir that reliably causes recurrent viremia upon ART cessation. Improved understanding of the mechanisms of HIV-mediated cell death could inform innovations designed to clear the latent reservoir. “Death Induced by Survival gene Elimination” (DISE) is an RNA interference (RNAi)-based mechanism that kills cells through short (s)RNAs with toxic 6mer seeds (pos. 2-7 of sRNA). These toxic seeds target reverse complementary seed matches in the 3’UTR of mRNA transcripts to decrease expression of hundreds of genes that are critical for cell survival. In most cells under normal conditions, highly expressed cell-encoded non-toxic microRNAs (miRNAs) block access of toxic sRNAs to the RNA-induced silencing complex (RISC) that mediates RNAi, promoting cell survival. We now report that infection of cells with HIV results in RISC-loading of an HIV-encoded miRNA, v-miRNA HIV-miR-TAR-3p, which kills cells by DISE through a noncanonical (pos. 3-8) 6mer seed. In addition, cellular RISC bound sRNAs shift to lower seed viability. Both these effects also occur with latent HIV provirus reactivation in J-Lat cells, a well-established cell model of HIV latency. Cells lacking expression of miRNA biogenesis genes Drosha, Dicer and Exportin 5 cannot produce protective miRNAs and therefore do not block RISC loading of the v-miRNA HIV-miR-TAR-3p. These mutant cells, as well as cells lacking expression of the RISC component Ago2, are hypersensitive to cell death via DISE induced by HIV infection. More precise targeting of the balance between protective and cytotoxic sRNAs could specifically and transiently increase silencing of cell survival genes to increase DISE. This could be a new addition to a “shock and kill” strategy to enhance depletion of the provirus reservoir during suppressive ART.

## Introduction

Active replication of HIV-1 (HIV) results in progressive loss of CD4+ T cells and a subsequent inability of the body to combat opportunistic infections (Acquired Immunodeficiency Syndrome, or AIDS). Current combination antiretroviral therapy (ART) inhibits viral replication and rescues CD4+ T cell levels. However, ART is unable to clear the infection due to the persistence of the provirus in cellular reservoirs. Efforts to clear the persistent provirus reservoir through forced reactivation of silent proviruses and induction of viral or immune mediated cytopathicity of the virus-producing cells (*i.e*., “shock-and-kill” strategies) have been unsuccessful thus far. This is partly due to resistance of CD4+ T cells that reactivate HIV to virus-induced cell death and to CD8+ T cell-mediated killing (1-5). Some mechanisms of resistance to immune cell-mediated killing have been suggested (6-9), but mechanisms why these cells are resistant to death mediated by HIV replication/antigen expression remain under study. Several mechanisms by which initial HIV infection is cytotoxic to infected and/or bystander cells have been reported (10-15) involving many viral genes (16-18) and various forms of cell death (19-24). Characterizing the mechanisms underlying the long-term survival of certain T cells that become persistent provirus reservoirs is critical to developing a cure.

RNA interference (RNAi) is a form of post-transcriptional regulation exerted by 19-22 nt long double-stranded (ds) RNAs that negatively regulate gene expression at the mRNA level. One of the two strands of the short dsRNA, the active guide strand, incorporates into the RNA induced silencing complex (RISC) (25) and the other strand (the inactive passenger strand) is degraded (26). RNAi is used as an endogenous form of gene regulation largely through the expression of microRNAs (miRNAs), short noncoding RNAs that coordinate silencing across hundreds of gene targets. This form of silencing is based on only a very short region of complete complementarity, the ‘seed’, at positions 2-7/8 of the guide strand (27, 28). Matching complementary regions (seed matches) predominantly located in the 3’UTR of mRNAs are targeted (29, 30) resulting in gene silencing (31). This targeting can be initiated with as little as a six nucleotide base-pairing between a guide RNA’s 6mer seed and the targeted mRNA (27, 28). miRNA biogenesis begins in the nucleus with transcription of a primary miRNA precursor (32). They are first processed by the Drosha/DGCR8 microprocessor complex into pre-miRNAs (33), which are then exported from the nucleus to the cytoplasm by exportin 5 (XPO5) (34). Once in the cytoplasm, Dicer/TRBP processes them further (35, 36) and these mature dsRNA duplexes are then loaded onto Argonaute (Ago) proteins to form the RISC (25). Consequently, the deletion/downregulation of either Drosha, Dicer, or XPO5 results in a general loss/reduction of miRNAs (37, 38).

We recently discovered death induced by survival gene elimination (DISE), a novel, tissue-independent, RNAi-based mechanism of cell death whereby toxic short (s)RNAs containing 6mer seed sequences (exerting 6mer seed toxicity) targeting essential survival genes are loaded into RISC complexes, resulting in inescapable cell death (38, 39). This kill code is likely more than 800 million years old (38). 6mer seed toxicity includes hallmarks of many different cell death pathways, including accumulation of DNA damage, activation of caspases, and mitochondrial stress (40). We previously tested 215 miRNAs coded for by 17 human pathogenic viruses including HIV (41) and provided evidence that many virus encoded miRNAs (v-miRNAs) have the capacity to kill cells through 6mer seed toxicity (41). Many of them were predicted to target mRNAs essential for cell survival using a noncanonical 6mer seed located at position 3-8 of the RISC-bound sRNA (R-sRNA). Therefore, we hypothesized that increasing DISE may counter a major problem with “shock and kill” strategies for HIV cure: the lack of cell death after reactivation of proviruses in reservoir cells under cover of ART.

HIV has been shown to code for 4 v-miRNAs (as deposited in miRBase 22.1). hiv1-miR-N367 has been shown to suppress the HIV gene nef (42). hiv1-miR-H1 can suppress cellular coding genes and miRNAs involved in regulation of apoptosis (i.e. Bcl-2) and RNAi (i.e. Dicer) (43). Finally, the HIV TAR element can give rise to two mature miRNAs: hiv1-miR-TAR-3p and hiv1-miR-TAR-5p (44). These miRNAs were found to be loaded into the RISC and miR-TAR-3p targets a number of apoptosis-regulating host genes including caspase-8 (45). Of note, caspase 8 is also suppressed in J-Lat cells, a derivative of Jurkat T cells that carry a transcriptionally latent HIV provirus and are often used to study latency reactivation (46). The v-miRNAs could therefore be important for cell survival and provirus persistence.

We now report that infection with HIV results in a shift of R-sRNAs from nontoxic to toxic, a phenomenon we recently described to contribute to therapy sensitivity of ovarian cancer (47). This is also aided by expression and uptake into the RISC of the HIV encoded v-miRNA miR-TAR-3p, which we determined to be much more toxic through its use of a noncanonical 3-8 6mer seed. Importantly, depleting cells of miRNAs renders them much more sensitive to HIV induced death. This was observed with deletion of either of the miRNA biogenesis genes Drosha, Dicer, or XPO5, but also with deletion of the RISC component Ago2. These findings add to characterization of mechanisms of HIV-induced cell death with implications for future improvement of “shock and kill” approaches to an HIV cure during suppressive ART via DISE.

## Results

### VSV-g pseudotyped HIV kills HCT116 cells that lack expression of genes required to generate mature miRNAs more effectively

We previously discovered a kill code embedded in miRNAs that can kill all cells by targeting genes critical for survival through a 6mer seed. Arrayed viability screens of all possible 4096 6mer seeds in a double stranded neutral siRNA backbone on three human and three mouse cell lines identified the most toxic 6mer seeds as being G rich (38, 48). G rich 6mer seeds were found to be present in a number of tumor suppressive miRNAs including miR-34a,c/449-5p and miR-15a,b/16-5p and most of their cytotoxic activity could be attributed to 6mer seed toxicity (38, 48). We provided evidence that the kill code is ancient (38) suggesting that viruses may also have adopted it. Subsequently, we discovered that human Kaposi sarcoma-associated herpes virus encoded v-miRNA miR-K12-6-5p acts as an oncoviral mimic through a 6mer seed shared with miR-15/16-5p (41). Interestingly, this shared seed match was not located at position 2-7, but at position 3-8 of the guide stand of the mature v-miRNA. This was validated by comparing the targeting of the mature miRNA with that of the neutral siRNA backbone used in the viability screens containing exclusively either the canonical (2-7) or the noncanonical (3-8) 6mer seed. Comparing the DISE inducing activity of 215 v-miRNAs with that of their 6mer seed containing siRNAs revealed that many v-miRNAs have the potential to kill cells by DISE possibly using the noncanonical 3-8 seed (41). This analysis included the four mature v-miRNAs encoded by HIV deposited at miRBase. When comparing the toxicity of the overexpressed HIV v-miRNAs (**Fig. 1A**, x-axis) with that of the siRNAs just containing the 6mer seed of that miRNA (**Fig. 1A**, Y axis), the HIV coded v-miRNAs showed a tendency to use the 3-8 seed (analysis based on the updated screening data on three human cell lines, 6merdb.org). In addition to direct cytotoxic effects of viral RNAs on infected cells, infection with HIV may also shift the ratio of cellular R-sRNAs to more toxic seeds essentially priming the cells to undergo DISE. This effect was recently found in ovarian cancer cells sensitive to chemotherapy compared to treatment resistant cancer cells (47) and in brain cells from Alzheimer’s disease patients compared to normal controls (49).

**Figure 1:**
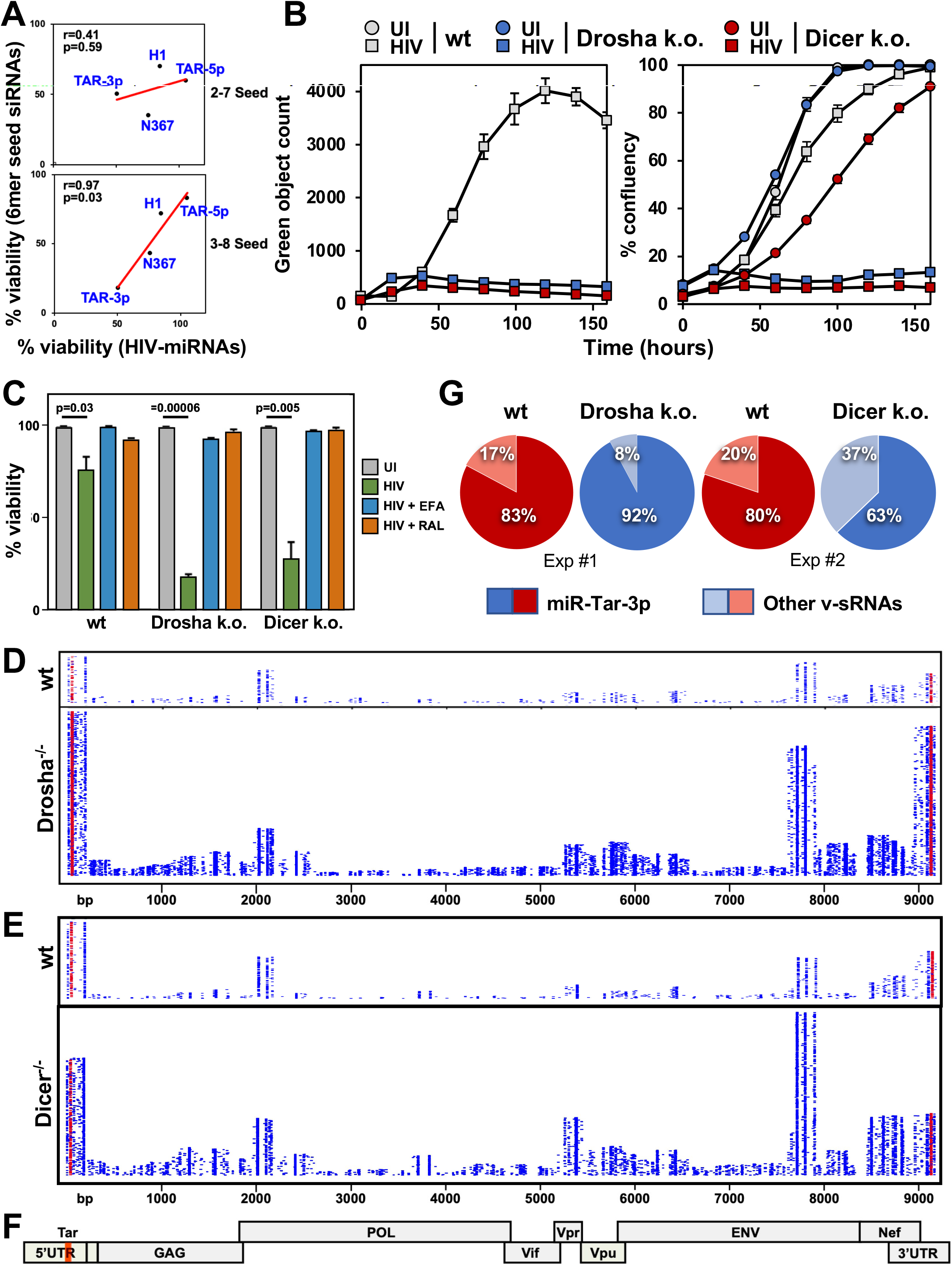
HIV kills HCT116 cells lacking either Drosha or Dicer more effectively. (**A**) Regression analysis showing correlation between the toxicity of the four putative HIV miRNAs (X-axis) and four siRNA duplexes carrying only matching 6mer seeds in a neutral backbone (Y-axis) as tested previously in three human cell lines (38, 48). Both the data for the canonical (top) and noncanonical 6mer seed (bottom) are plotted. P-values were calculated using Pearson correlation analysis. (**B**) Change in green object count (HIV infected cells) (left) and confluency (right) over time of HCT116 wt, Drosha k.o., or Dicer k.o. cells. Cells were infected with VSV-G pseudotyped HIV R9 ΔEnv iGFP virus (175 ng p24/ml) (50). UI, uninfected. (**C**) Wt, Drosha k.o., or Dicer k.o. HCT116 cells were infected with VSV-G pseudotyped HIV R9 ΔEnv iGFP virus (175 ng p24/ml) in the presence of either 10 μM of efavirenz (EFA) or 10 μM of raltegravir (RAL), as indicated. 72 hours after challenge, cell-death was quantified. Student’s t-test p-values are shown. (**D, E**) Whole HIV genome alignment of reads pulled down with Ago proteins in wt and Drosha k.o. (D) or in wt and Dicer k.o. (E) HCT116 HIV infected as in B. Each horizontal line represents one read. Reads from both replicates per condition are displayed. miR-TAR-3p derived reads are shown in red. (**G**) Percent HIV derived reads from miR-TAR-3p in the two experiments shown in D and E. (**F**) Schematic of the HIV-1 genome. The location of the TAR loop is indicated in orange. Note: the ΔEnv virus still expresses robust amounts of ENV derived RNA (see reads aligned to the ENV gene) as it carries a complete ENV gene with both a KDEL ER-retention signal and a stop codon included (87). While the reads from the TAR loop aligned at both the 5’ and 3’ LTR, it was recently shown that the miR-TAR derived reads are derived from the 5’ TAR loop (88).

To test whether the toxicity exerted by HIV infection had a component of 6mer seed toxicity, we infected wild-type (wt), Drosha knock-out (k.o.), and Dicer k.o. HCT116 cells with a VSV-G pseudotyped HIV R9 ΔEnv iGFP reporter virus (**Fig. S1A**) described previously (50). We chose HCT116 cells for these experiments as they provide a standardized way to test whether a cell death stimulus has a contribution of 6mer seed toxicity as used previously (38, 39, 51). Both Drosha and Dicer k.o. cells lack most canonical miRNAs containing nontoxic 6mer seeds protecting cells from DISE (52). Hence, they are hypersensitive to DISE inducing stimuli. Tracking virus infected (GFP+) cells in an IncuCyte Zoom microscope over 7 days showed significant reduction in infection of the two mutant cell lines when compared to wt cells (**Fig. 1B**, left graph). This was likely due to the observation that HIV infected Drosha and Dicer k.o. cells succumbed to cell death much more quickly than wt cells (**Fig. 1B**, right graph). This observation was further confirmed in a cell death assay (**Fig. 1C**). A similar observation was made with HCT116 XPO5 deficient cells which are also defective in producing most mature miRNAs (37); two XPO5 k.o. clones tested were also hypersensitive to HIV induced cell death (**Fig. S1A** and **S1B**). VSV-G pseudotyped HIV enters the cells through the LDL-R receptor (53) which could be regulated by cellular (c-)miRNAs and hence in the absence of these miRNAs in the mutant cells, LDL-R could be upregulated potentially explaining the increased effect of HIV on these cells. However, flow cytometry analysis showed that none of the mutant cells had upregulated surface LDL-R. On the contrary, all the mutant cells showed reduced LDL-R relative to wt cells (**Fig. S1C**).

To test whether the HIV induced cytotoxicity is due to early viral replication events, we treated cells with either the RT inhibitor efavirenz (EFA) or the integrase inhibitor raltegravir (RAL). Cells were infected as before and cell viability was monitored 3 days after infection (**Fig. 1C**). Both inhibitors completely prevented HIV induced cell death. This indicates that post-integration events, rather than the viral RNA, RNA-DNA hybrid, or reverse transcribed DNA cause enhanced death of Drosha/Dicer k.o. cells. These results are consistent with a model wherein HIV induces cell death through either its gene products, RNAs produced by the virus, or cellular changes in the infected cells. The fact that cells devoid of genes required for the generation of canonical miRNAs were hypersensitive to this form of cell death suggested that other sRNAs, either virus or host cell derived, could be involved in cell survival.

To directly determine whether HIV induced toxicity in the cells with decreased miRNA biogenesis involved changes in R-sRNAs, we performed an Ago1-4-RNA precipitation combined with high-throughput sequencing (Ago-RP-Seq) of wt, Drosha and Dicer k.o. HCT116 cells infected with VSV-G pseudotyped HIV as described (38, 47, 51). Infection with HIV did not have a major effect on the total miRNA content in either wt or the two k.o. cells (**Fig. S2A**). As expected, the two k.o. cells had a very low miRNA content and many of the R-sRNAs were not c-miRNAs. Consistent with a previous analysis (54), only a few HIV derived reads were found in the RISC of wt cells (**Fig. 1D, S2B**). In contrast, both Dicer and Drosha k.o. HCT116 cells took up >5-10 times more HIV reads into their RISC (**Fig. 1D, 1E, S2B**).

Multiple HIV derived miRNAs have been described (43, 45, 54, 55). In a comprehensive analysis of HIV-derived RNAs (56), the sequences of reads derived from the HIV TAR RNA element, which is the source of the putative HIV-derived miRNA miR-TAR, were provided (54) (**Fig. S3A**). TAR derived reads were derived from both arms of the TAR hairpin (**Fig. S3B**). In the hairpin, the two read clusters detected are juxtaposed with 3’ overhangs, which suggested processing by Dicer and loading into the RISC (57, 58) and consequently miR-TAR processing has been shown to be dependent on Dicer (44). The majority of the HIV derived R-sRNAs in our analyses aligned with the HIV UTR and the TAR loop, the location of the putative v-miRNA miR-TAR (45) (**Fig. 1D** and **1E**). Interestingly, while they constitute a smaller fraction of all HIV derived R-sRNAs when compared to the Drosha k.o. cells, reads derived from the processed mature v-miRNA miR-TAR-3p were still RISC loaded in larger quantities in cells completely deficient of Dicer (37) (**Fig. 1G**) suggesting that they can be generated without the involvement of Dicer.

### HIV infection results in an increase in R-sRNAs with toxic 6mer seeds

We previously reported that the ratio of R-sRNAs with toxic versus nontoxic 6mer seeds can prime cells for DISE (47). We therefore tested whether the infection with HIV influenced the average seed toxicity of either all R-sRNAs or virally derived R-sRNAs. We used SPOROS, a recently developed bioinformatics pipeline that allows analysis of RNAseq data in a gene agnostic way solely based on the predicted 6mer seed toxicity of each read (59). One of the SPOROS outputs is a “seed tox” graph which plots the read counts of all sRNAs (Y axis) against the predicted cell viability based on our toxicity screen of all 4096 6mer seeds (X-axis) (38, 48). We analyzed two independent experiments, one comparing wt cells with Drosha k.o. cells (**Fig. 2A**) and the other comparing wt cells with Dicer k.o. cells (**Fig. 2B**). As reported previously, in wt cells >95% of sRNAs were miRNAs with mostly nontoxic seeds (**Fig. 2A, B**, top left graph). In contrast, many sRNAs other than miRNAs (e.g., tRNA fragments) entered the RISC (labeled in black) in Dicer k.o. (**Fig. 2B**, top right graph), and to a greater extent in Drosha k.o. cells (**Fig. 2A**, top right graph).

**Figure 2:**
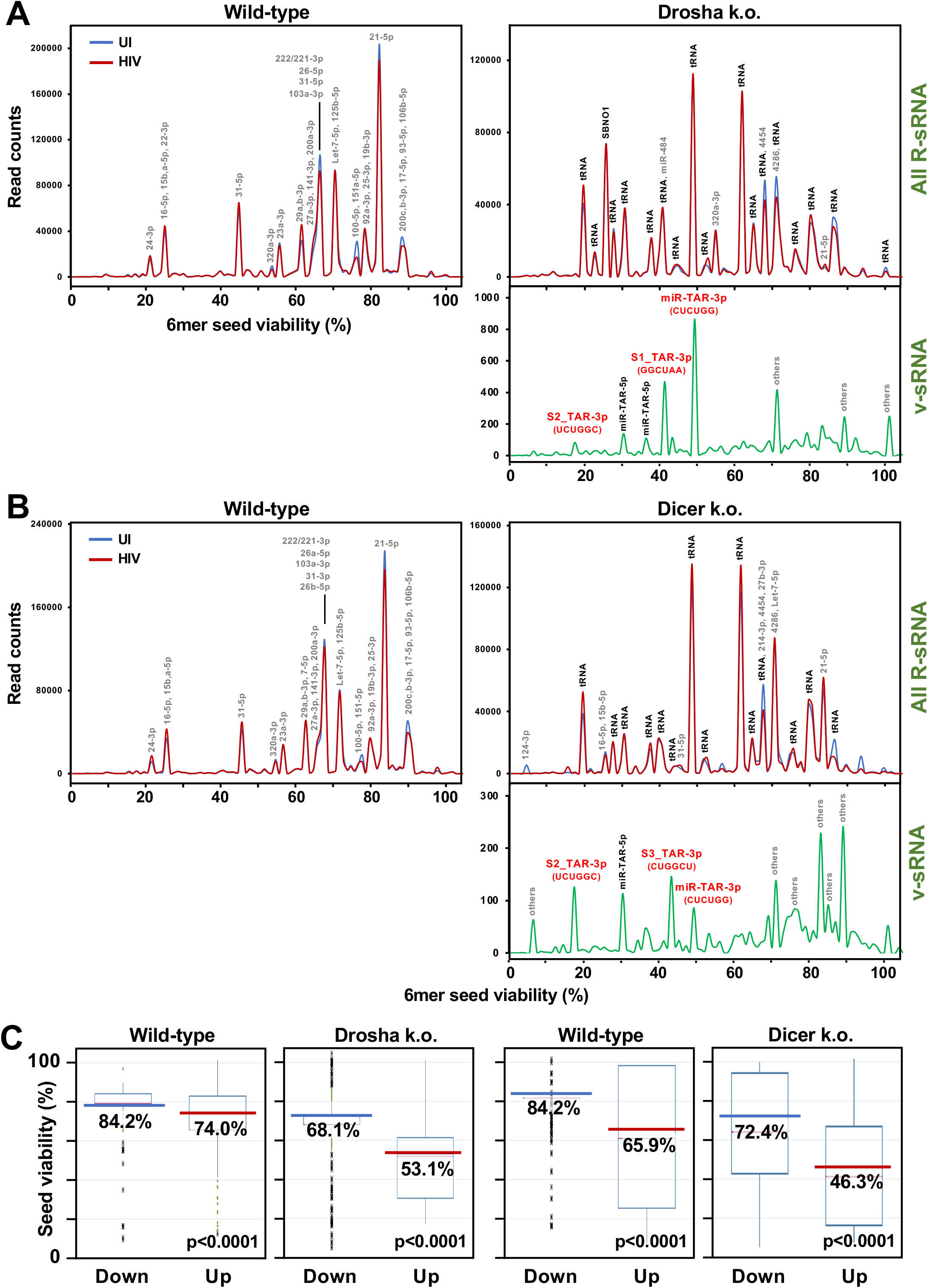
HIV infection results in displacement of nontoxic cellular R-sRNAs with toxic c-miRNAs and highly toxic v-miRNAs. (**A, B**) Seed toxicity graphs of all R-sRNAs in wt and Drosha k.o. cells (A) or wt and Dicer k.o. cells (B) uninfected or 28 hours after infection with VSV-G pseudotyped HIV. miRNAs that contribute to peaks with >5,000 reads are labeled. For each labeled peak, miRNAs are listed in the order of abundance. For both experiments a seed toxicity graph is shown with only the v-miRNAs (in green). Three miR-TAR-3p isomiRs with their respective 2-7 seed are labeled in red. (**C**) Seed toxicity box plots showing the median seed viability (%) of R-sRNAs significantly up or downregulated (>1.5 x, p< 0.05) in wt, Drosha k.o. or Dicer k.o. cells after HIV infection. The median is highlighted.

Most of the HIV derived R-sRNAs in Drosha k.o. cells were from miR-TAR with the most abundant ones from the 3p arm of this v-miRNA (**Fig. 2A**, right bottom graph). Interestingly, three different species (isomiRs) were detected (labeled in red as miR-TAR-3p, S1_TAR-3p and S2_TAR-3p). While the number of miR-TAR derived reads was relatively small, we recently showed that very small numbers of CD95L derived R-sRNAs with toxic seeds could kill HeyA8 ovarian cancer cells through RNAi (51). Interestingly, the relative amounts of miR-TAR-3p isomiRs detected in the RISC of Dicer k.o. cells (**Fig. 2B**, bottom right graph) were different from the ones found in Drosha k.o. cells (**Fig. 2A**, top right graph) suggesting that a nuclease other than Dicer could be generating these reads.

To determine overall changes in R-sRNAs with respect to their seed viability, we plotted the median seed viability of all R-sRNAs (a SPOROS output) in the different genotypes (**Fig. 2C**). HIV infection caused a general enrichment of R-sRNAs carrying toxic seeds versus nontoxic seeds (**Fig. 2C)**. This was mainly due to a shift in abundant cellular R-sRNAs to more toxic ones. These findings were confirmed in an independent set of experiments directly comparing the response of wt, Drosha and Dicer k.o. cells (**Fig. S4**). In summary, these data are consistent with previous reports of HIV infection suppressing expression of c-miRNAs that we determined to carry nontoxic 6mer seeds (60, 61). At the same time, virus-derived sRNAs appeared in the RISC (mostly miR-TAR-3p). Our data indicate that HIV infection can shift the balance from nontoxic to more toxic cellular R-sRNAs.

### HIV-miR-TAR-3p contains a highly toxic noncanonical 6mer seed

In our analysis of HIV-derived R-sRNAs in Drosha k.o. cells, we found three different miR-TAR-3p isomiRs with different start sites and hence different predicted 6mer seeds (**Fig. 3A**). For the two main isomiRs, the miRBase annotated version and version S1, the noncanonical 6mer seed (pos. 3-8) was predicted to be more toxic than that canonical seed (pos. 2-7). Toxicity of the isomiRs was established by transfecting them at 1 nM into cells (**Fig. 3B-D**). As expected, both Dicer and Drosha k.o. cells with reduced amounts of c-miRNAs were more susceptible to this toxicity than wt cells. Toxicity stemmed from the 6mer seed as shown by transfecting wt or Drosha k.o. cells with siRNAs that only carried the different 2-7 or the more toxic 3-8 seeds of the two main isomiRs (**Fig. 3E, F**). These data suggested that the main HIV derived reads in the RISC of infected HCT116 cells were from the HIV miRNA miR-TAR-3p which may therefore contribute to the cytotoxicity of HIV through 6mer seed toxicity. This was also consistent with the observation that cells which lack most canonical c-miRNAs were hypersensitive to cell death induced by transfection with small amounts of miR-TAR-3p implicating the mechanism of 6mer seed toxicity in HIV infected cells as well.

**Figure 3:**
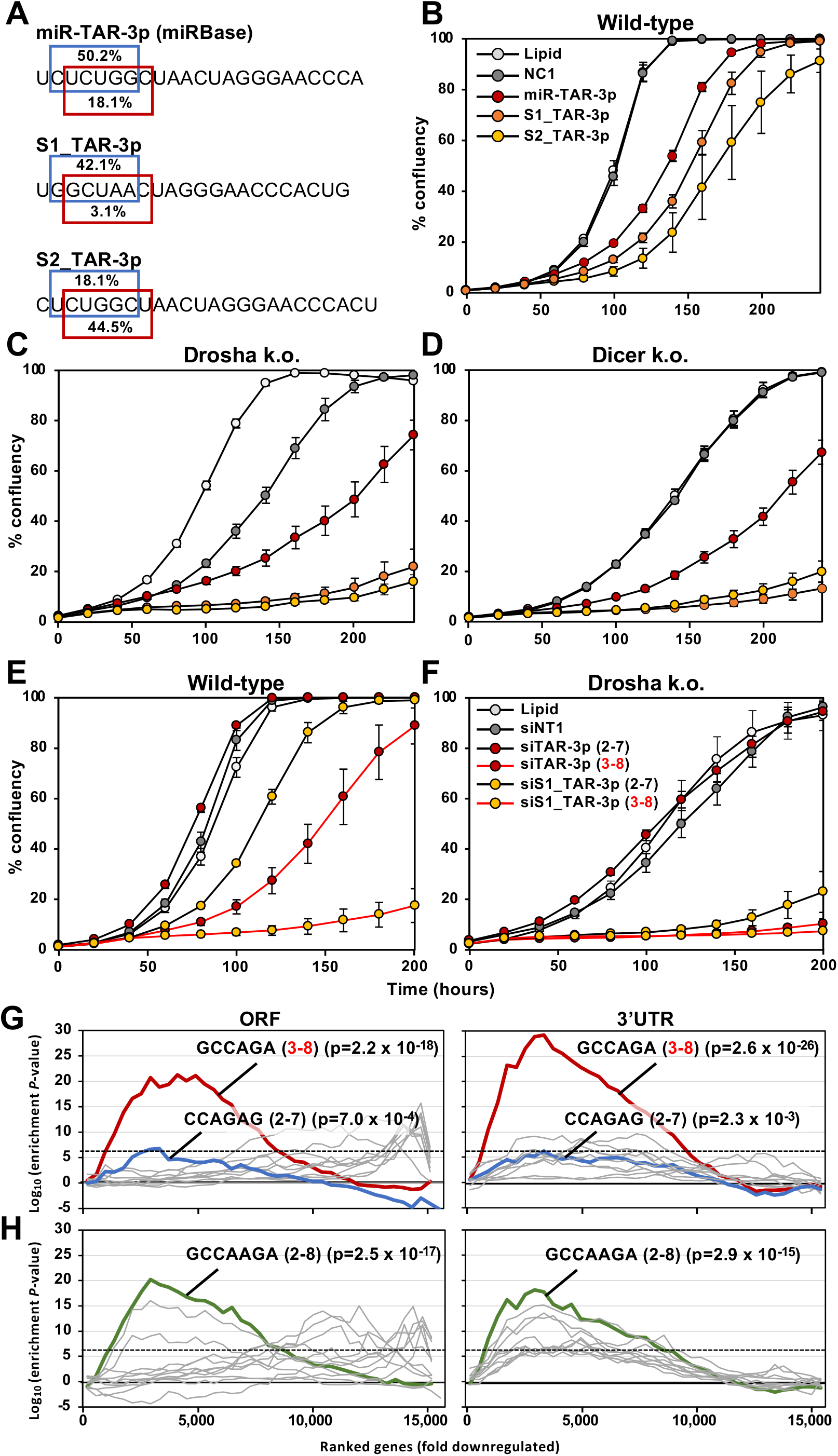
HIV-miR-TAR-3p isomiRs contain toxic 6mer seeds and can kill cells through DISE. (**A**) Three miR-TAR-3p isomiRs detected in the RISC of HIV infected cells. *Top*, the canonical miR-TAR-3p as deposited in miRbase. *Center*, S1_miR-TAR-3p, an isomiR with 5’ end 4 nucleotides downstream from the canonical start site. *Bottom*, S2_miR-TAR-3p, an isomiR with 5’ end 1 nucleotide downstream from the canonical start site. In each case, the predicted seed viability of the canonical 2-7 (blue) and the noncanonical 3-8 (red) 6mer seed is indicated. (**B**-**D**) Confluency over time of HCT116 wt (B), Drosha k.o. (C), or Dicer k.o. (D) cells treated with either lipid alone, or transfected with 1 nM of control miRNA NC1, HIV-miR-TAR-3p, S1_miR-TAR-3p, or S2_miR-TAR-3p. Samples were analyzed in quadruplicates and SE is shown. (**E, F**) Confluency over time of HCT116 wt (E) or Drosha k.o. cells (F) treated with either lipid alone, or transfected with 1 nM of either siNT1, siTAR-3p (2-7 seed), siTAR-3p (3-8 seed), siS1_TAR-3p (2-7 seed), or siS1_TAR-3p (3-8 seed). Samples were analyzed in quadruplicates and SE is shown. (**G, H**) Sylamer analysis (6mers in G, 7mers in H)) for the list of open reading frames (ORFs; left) and 3’UTRs (right) of mRNAs in cells treated with 10 nM miR-TAR-3p for 24 hrs sorted from down-regulated to up-regulated. Curves for the 10 most enriched seeds are shown. Enrichment of the GCCAGA sequence targeted by miR-TAR-3p through the noncanonical 3-8 6mer seed is shown in red (G), the CCAGAG sequence targeted by miR-TAR-3p through the canonical 2-7 6mer seed is shown in blue (G), the GCCAGAG sequence targeted by miR-TAR-3p through the canonical 2-8 7mer seed is shown in green (H). The stippled line corresponds to a p-value threshold of 0.05 after Bonferroni correction for the number of words tested (4096). Bonferroni-adjusted p-values are shown.

Our comparison of the toxicity exerted by the four HIV miRNAs with the toxicity of just their 6mer seeds embedded in a neutral siRNA backbone suggested that HIV miRNAs may target through a noncanonical 6mer seed in position 3-8 (**Fig. 1A**). This was consistent with our analysis of 215 v-miRNAs that suggested that many v-miRNAs may use this noncanonical seed (41). Because the two main species of miR-TAR-3p we detected were predicted to be much more toxic when targeting through the 3-8 6mer seed, we were wondering whether introduction of the authentic v-miRNA would target through a canonical 2-7 or a noncanonical 3-8 seed. To test this, we transfected HCT116 cells with 10 nM of either a control miRNA (NC1) or miR-TAR-3p and 24 hours after transfection subjected the RNA to an RNAseq analysis. Enrichment of targeted 6mer seed matches in the downregulated genes was determined using a Sylamer analysis (62). This analysis detected a very strong enrichment of the noncanonical seed match GCCAGA in the 3’UTR of downregulated genes (red line in **Fig. 3G**, right panel) and a much lower enrichment of the canonical CCAGAG seed match. A similar but less pronounced enrichment was also found in the ORF. Enrichment of the canonical 7mer seed match was also found, but was less pronounced (**Fig. 3H**, green lines). These data suggest that miR-TAR-3p targets mostly through its much more toxic noncanonical 3-8 6mer seed even more than with its canonical 7mer seed. In contrast to most c-miRNAs, there was substantial targeting of miR-TAR-3p in both the 3’UTR and the ORF. In summary, even though the number of miR-TAR-3p derived R-sRNAs is low, they are expected to have very high toxicity through 6mer seed toxicity and a noncanonical targeting.

### Deletion of Drosha in Jurkat T cells renders them more sensitive to HIV induced cytotoxicity

To determine whether the lack of most miRNAs and/or Drosha expression could also affect susceptibility of T cell derived lines to HIV-induced cell death, we generated and single cell cloned Drosha k.o. Jurkat T cells (**Fig. 4A**). Two clones were selected (E11 and F6) that had substantially reduced Drosha protein expression. Both clones had a significantly reduced ability to produce mature miR-21 and miR-200b, but not miR-320 as detected by qPCR (**Fig. S5A**); the latter is generated independent of Drosha (37). The reduction in mature miRNAs was not as pronounced as in Drosha k.o. HCT116 cells (37) presumably due the fact that we could not generate Jurkat clones that were completely devoid of Drosha given its requirement for cell survival in this cell line (see methods). Both Drosha k.o. Jurkat clones experienced a greater loss of viability following challenge with an VSV-G pseudotyped HIV ΔEnv iGFP reporter virus than the wt cells (**Fig. 4B**), suggesting that the extent of Drosha knockdown was sufficient to see the effect of HIV cytotoxicity as observed before in the HCT116 cells. As in the HCT116 cells, this effect was not due to an increased expression of the LDL-R entry receptor on the surface of the two k.o. clones and LDL-R expression was instead decreased in these clones (**Fig. S5B**). As seen with the results in HCT116 cells, seed toxicity plots of all R-sRNAs showed that in both Drosha k.o. clones 24 hours after infection with HIV, seed viability of R-sRNAs shifted to more toxic seeds (**Fig. 4C**). Wt cells did not show this effect and were not killed by HIV at the concentration used, when tested for viability after 72 hours (as shown in **Fig. 4B**).

**Figure 4:**
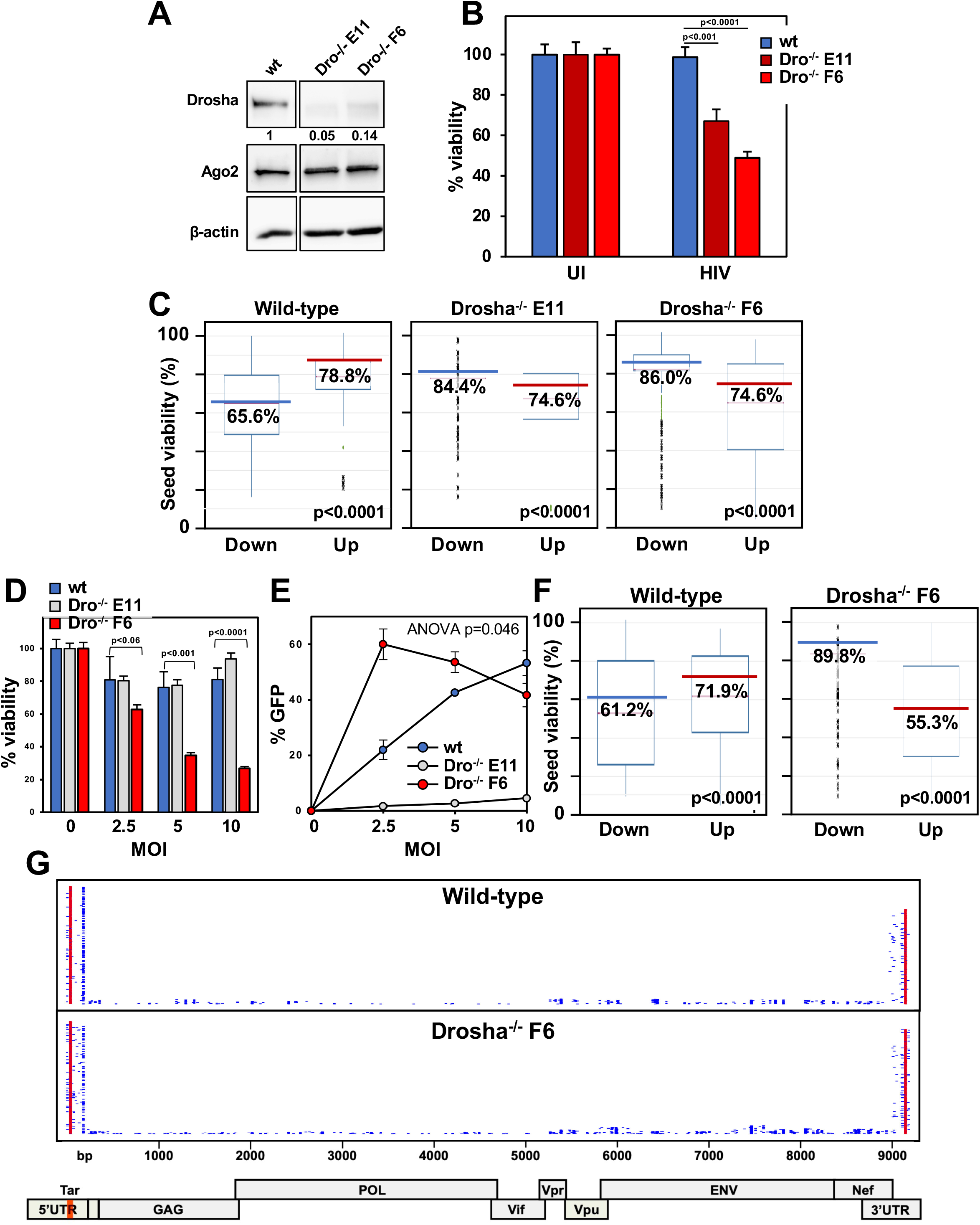
HIV kills Jurkat T cells lacking Drosha more effectively. (**A**) Western blot analysis of wt Jurkat T cells or two Jurkat single cell clones deficient in Drosha. Protein expression was quantified by densitometry relative to Actin. (**B**) Cell viability of wt Jurkat cells, wt or Drosha k.o. clones determined 72 hours after infection with 25 ng/ml VSV-G pseudotyped HIV. (**C**) Seed toxicity box plots showing the median seed viability of R-sRNAs significantly enriched (up) or depleted (down) in HIV infected Jurkat cells and clones. The median is highlighted. (**D**) Cell viability of wt Jurkat cells or two Drosha k.o. clones determined 8 days post-infection with replication competent HIV NL4-3 Nef:IRES:GFP virus tested at MOI 2.5, 5 and 10. (**E**) GFP positivity of the cells in (D) infected with replication competent HIV NL4-3 Nef:IRES:GFP virus tested at MOI 2.5, 5 and 10, 8 days post-infection. (**F**) Seed toxicity box plots showing the median seed toxicity of RISC R-sRNAs significantly enriched or depleted in replication competent HIV infected wt Jurkat cells (left) or Drosha k.o. clone F6 (right). The median is highlighted. (**G**) Whole HIV genome alignment of reads pulled down with Ago proteins in wt Jurkat cells (*top*) or Drosha k.o. clone F6 (*center*). Each horizontal line represents one read. Reads from replicates per condition are displayed. miR-TAR-3p derived reads are shown in red. *Bottom*: schematic of the HIV-1 genome. The location of the TAR loop is indicated in orange. Student T-test (B, D) or ANOVA p-values (E) are shown. P values of deregulated seed viabilities (C, F) were determined using a Kruskal–Wallis median rank test.

To test whether this was also found in a T cell line infected with T cell tropic replication proficient HIV, we infected wt Jurkat and the two Drosha k.o. clones with HIV NL4-3 Nef:IRES:GFP virus. Similar to data with the HCT116 cells, one Drosha k.o. clone (F6) was significantly more susceptible to HIV induced cell death (**Fig. 4D, E**). Clone E11 was not efficiently infected by HIV, and this was likely because it had substantially reduced CD4 surface expression (**Fig. S5C**). In contrast, clone F6 had similar expression of both entry receptors for HIV infection – CXCR4 and CD4. Consistent with the level of cell death induction in clone F6, R-sRNAs enriched in infected cells contained substantially more toxic seeds than the R-sRNAs in infected wt cells (**Fig. 4F**). In addition, HIV derived R-sRNAs were predominantly comprised of miR-TAR-3p reads (**Fig. 4G**) as seen with the HCT116 cells.

### HIV reactivation causes a shift of R-sRNAs to more toxic seeds in a model of HIV latency

Given our data indicating that HIV-induced cytotoxicity was related to post integration events, we next studied J-Lat cells, a derivative of Jurkat T cells that carry a transcriptionally latent HIV provirus and are often used to study latency reactivation (46). To release latent HIV, cells were treated with PMA to induce viral transcription. HIV induction caused profound cell death in these cells (**Fig. 5A**). The amount of RISC-bound v-miRNAs quadrupled in the PMA-induced cells (**Fig. 5B**) and specific changes in sRNAs occurred that resulted in a significant overall reduction in seed viability of all R-sRNAs (**Fig. 5C**) as well as a substantial enrichment of R-sRNAs with toxic seeds (**Fig. 5D**) despite the low absolute level of RISC-bound miR-TAR-3p.

**Figure 5:**
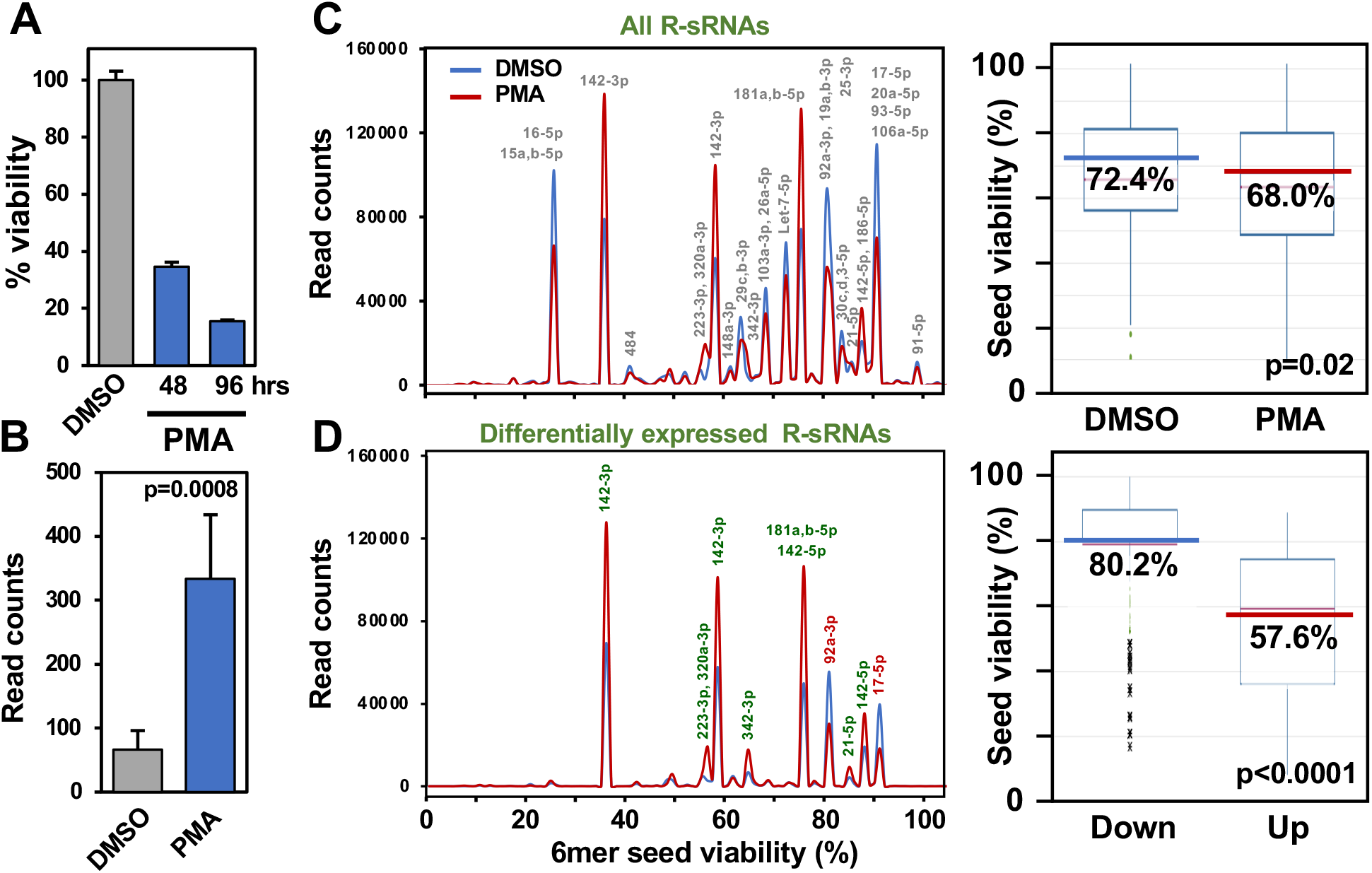
HIV reactivation causes a shift in toxicity of R-sRNAs in a model of HIV latency. (**A**) Viability of J-Lat cells determined 48 and 96 hours after addition of PMA. (**B**) Total HIV derived R-sRNAs in J-Lat cells treated with either DMSO control or PMA. Student’s t-test p-value is shown. (**C, D**) *Left panels*, seed toxicity graph of total (C) or only differentially expressed (D) R-sRNAs in J-Lat cells with latent or induced HIV. *Right panels*, seed toxicity box plots showing the median seed viabilities of all R-sRNAs (**C**) or only the ones enriched or depleted in J-Lat cells after induction of virus (D). The median is highlighted. P values of deregulated seed viabilities were determined using a Kruskal–Wallis median rank test.

### Deletion of Ago-2 expression increases susceptibility of cells to HIV induced toxicity

To test the role of the functionally essential RISC component Ago2 in HIV induced cytotoxicity, we again used the only cells we have available that can tolerate a homozygous knockout of Argonaute proteins: HCT116 cells completely deficient in either Ago1 alone, Ago2 alone, or the triple knock out (TKO) of Ago1,2,3. Each of the Ago-deficient cells were infected with HIV (**Fig. S6A-E**). At the chosen HIV titer, we did not see much of an effect of infection on cell death in wt cells (**Fig. S6A**). In contrast, both Ago2 and Ago1,2,3 TKO, but not Ago1 k.o., cells were hypersensitive to HIV induced toxicity (**Fig. S6B-D**). This increase in sensitivity was similar to what we observed in the Drosha k.o. cells (**Fig. S6E**).

There are four different Argonaute genes in the human genome. While Ago1,2,3 TKO cells were devoid of the three main Ago proteins, they could still express Ago4. In these cells, Ago4 was strongly increased both on the protein and the mRNA level (**Fig. S7A, B**). The contribution of Ago4 to the toxic effects of HIV could not be tested because Ago4 could not be knocked down in the TKO cells (**Fig. S7C**). This could either be due to the fact that at least one Ago protein needs to be expressed for RNAi to function or due to the observation that no cell, not even an embryonic stem cell, can survive after deletion of all four Ago genes (63). We concluded that Ago1 and 2 do not play a major role in the cytotoxicity of HIV. While Ago4 protein is highly upregulated in Ago1,2,3 TKO cells, whether it contributes to HIV cytopathicity cannot be fully assessed. In summary, our data suggest that the absence of either Drosha, Dicer, or Ago2 allowed HIV to induce cell death much more efficiently at least in part through the DISE mechanism however, a general antiviral activity of these genes cannot be excluded.

## Discussion

Multiple modes of HIV induced cell death have been previously proposed. Canonical apoptosis was the first cell death pathway that was implicated in the death of HIV infected and uninfected bystander cells through a number of stimuli, including: death ligands (FasL, TNF and TRAIL) expressed on immune cells (11); direct cytotoxicity of a number of soluble HIV proteins (e.g. Gp120, Tat, Nef, Vpr) (16-18); activation-induced cell death driven by a chronically activated and hyper-inflammatory immune state (10, 11); and through DNA-dependent protein kinase (DNA-PK) activation during viral integration (12). More recently, a series of studies (13-15) described how CD4+ T cells abortively infected with HIV die through caspase-1 dependent pyroptosis. This form of inflammatory cell death is thought be the primary mechanism by which uninfected bystander cells in the tissue die during infection (13). Other forms of cell death implicated to occur in HIV infected CD4+ T cells are autophagy (19, 20), necrosis (21, 22), necroptosis (23), and mitotic catastrophe (24).

DISE is a combination of various cell death pathways that are activated simultaneously due to targeting of multiple survival genes that abolish protection from different forms of cell death by sRNAs in a miRNA-like fashion. While each miRNA can often target hundreds of genes, the function of a miRNA is defined by the nature of the genes in its targeted network. We previously identified a kill code embedded in miRNAs that requires as little as 6 nucleotides (the 6mer seed) (38). When this 6mer seed is G rich, then any sRNA that contains this seed and when loaded into the RISC can kill cells through DISE by targeting C-rich seed matches located in the 3’UTR of genes whose expression are essential for cell survival (38, 48). We previously explored whether viruses could code for v-miRNAs carrying toxic seeds (41). We identified multiple v-miRNAs including HIV encoded v-RNAs as potentially being toxic to cells through 6mer seed toxicity. Interestingly, we found evidence of use of a noncanonical (pos. 3-8) 6mer seed for a number of v-miRNAs.

In the context of HIV infection, miRNAs have been shown to be relevant in two ways. HIV encodes at least four v-miRNAs (64). These have been shown to be involved in regulating viral replication, cell survival, host gene expression, and latency (45, 65, 66). However, the expression of these v-miRNAs are controversial (64). In addition, the expression of a number of c-miRNAs is affected by the virus and some have anti-viral activities (reviewed in (66)). Of the v-miRNAs, the only v-miRNA we detected in the RISC of infected cells in significant numbers was miR-TAR-3p. We determined that upon transfection, miR-TAR-3p gains toxicity by using the noncanonical (pos. 3-8) 6mer seed.

The discovery of DISE/6mer seed toxicity allows a novel way of looking at any sRNA that can be taken up by the RISC and exert RNAi. In addition to the action of individual miRNAs on specific targets regulating various cellular functions, 6mer seed toxicity requires assessment of the entirety of all R-sRNAs with respect to their seed toxicity and changes in the average seed viability to contribute to cell fate regulation. To study the potency of R-sRNAs to kill cells, we recently developed SPOROS, a bioinformatics pipeline that allows analysis of all RISC bound reads in a gene agnostic and standardized fashion (59). Using SPOROS, we now provide evidence that HIV infection causes a shift of R-sRNAs to higher toxicity. Our data suggest both an involvement of a v-miRNA as well as a contribution of cellular R-sRNAs to HIV induced cell death. Such a shift of cellular R-sRNAs to more toxic seeds may prime cells for DISE similar to other systems we have tested recently (47, 49).

It is intriguing that deletion of multiple genes involved in either the maturation (Drosha, Dicer or XPO5) or function of miRNAs (Ago2) caused an increase in susceptibility to 6mer seed toxicity. Particularly, the results in cells deficient in Ago2 (but not Ago1) expression may argue against a contribution of the RISC in HIV induced cytotoxicity. However, the data on Ago1,2,3 TKO cells does not exclude an involvement of the fourth Ago protein in the human genome, Ago4. Likely as a compensatory mechanism, Ago4 was found to be highly upregulated in the Ago1,2,3 TKO cells. While Ago4 was shown not to play a role in siRNA mediated activities, it can bind miRNAs and mediate their function (63). Unfortunately, we could not knockdown Ago4 to test its involvement as this requires the presence of Ago2 and knocking out Ago4 would also not be possible as any cell needs to express at least one Ago gene in order to survive (63). The contribution of miR-TAR-3p and its isomiRs to HIV induced toxicity via RISC function can therefore not be excluded at this point. Mechanistically, given the nature of 6mer seed toxicity which involves targeting a network of genes, it is difficult to perform rescue experiments. HIV has been shown to be cytopathic through the activity of the protein products of many of its genes, including Nef, Vpr and Tat (16-18). It is unlikely that 6mer seed toxicity is the only mechanism mediating HIV cytopathicity. It is therefore not to be expected that even eliminating all RISC activity would completely block HIV induced cytotoxicity.

Some c-miRNAs have been shown to have antiviral activities (reviewed in (66)) and that would suggest a higher expression of HIV in cells lacking miRNA biogenesis enzymes or Ago2. We did detect a higher expression of HIV in all mutant cells (data not shown) and this could be due to altered miRNA levels/activity affecting expression levels of the HIV entry receptors. However, we can exclude this effect a mode of action as we did not find an increased expression of any of the entry receptors of the HIV viruses we used in our study in the mutant cells. In addition, the substantial reduction in the seed viability of cellular R-sRNAs in J-Lat cells after induction of latent HIV expression suggests a substantial contribution of 6mer seed toxicity to HIV induced cell death. Finally, the much higher sensitivity of Drosha or Dicer k.o. cells to the toxic effects of miR-TAR-3p or its seed containing siRNAs also supports the increase in sensitivity of cells lacking c-miRNAs to the DISE inducing properties of the v-miRNA.

Interestingly, Drosha, Dicer, XPO5, and Ago2 (but not Ago1) also function in the DNA damage response. Both Dicer and Drosha facilitate DNA repair through the generation of Dicer and Drosha-dependent small RNAs (DDRNAs) (67), which mediate recruitment of repair complexes to the sites of DNA damage (68). Drosha also promotes non-homologous end joining by interacting with RAD50 (69). XPO5 has been reported to promote genomic stability by exporting pre-miRNAs from the nucleus that facilitate DNA repair (70). Finally, Ago2 is indispensable for its function in Rad51 recruitment and homologous recombination repair (71). HIV has been linked to DNA damage through its integration into the genome (72) and RNA viruses in general are known to interfere with the DNA damage response through multiple mechanisms (73).

In summary, our data suggests that HIV infection causes a shift of mostly cellular R-sRNA to more toxic seeds. This could prime cells to DISE, which could be enhanced by the v-miRNA miR-TAR-3p which carries a toxic 6mer seed and may even gain higher toxicity through a noncanonical 3-8 seed. Our data provides multiple new avenues to explore novel cell death mechanisms that could be used to kill latent HIV.

## Methods

### Cell lines

The human colorectal cancer cell line HCT116 and its mutant variants Drosha k.o. #40, Dicer k.o. #43, Xpo5 k.o. #19 and 19-1 (37) were obtained from the Korean Collection for Type Cultures (KCTC) and cultured in McCoy’s 5A medium (ATCC #30-2007) supplemented with 10% heat inactivated FBS (Sigma-Aldrich #14009C) and 1% penicillin/streptomycin (Corning #30-002-CI). HCT116 Ago2 k.o. (a kind gift from Joshua T. Mendell, UT Southwestern) (74), Ago1 k.o. and Ago123 k.o. (kindly provided by David Corey, UT Southwestern) (75) were cultured in McCoy’s 5A medium (ATCC #30-2007) supplemented with 10% FBS and 1% penicillin/streptomycin. The human embryonic kidney cell line 293T was obtained from ATCC (CRL-3216) and cultured in DMEM medium (Corning #10-013 CM), supplemented with 10% heat inactivated FBS (Sigma-Aldrich #14009C) and 1% penicillin/streptomycin (Corning #30-002-CI). The human T lymphoblast cell line Jurkat, Clone E6-1 was obtained from ATCC (TIB-152) and cultured in suspension in RPMI1640 medium (Corning #10–040 CM), supplemented with 10% heat inactivated FBS (Sigma-Aldrich #14009C) and 1% penicillin/streptomycin (Corning #30-002-CI). Jurkat-derived HIV-1 latency model cell line, J-Lat (kindly provided by Judd Hultquist, Northwestern University) was cultured in suspension in RPMI1640 medium (Corning #10–040 CM), supplemented with 10% heat inactivated FBS (Sigma-Aldrich #14009C), 1% penicillin/streptomycin (Corning #30-002-CI) and 10 μM antiretroviral cocktail. Jurkat Drosha k.o. cell pool was designed and engineered using the CRISPR-Cas9 system by Synthego. The sgRNA sequence: CACAGAAUGUCGUUCCACCC targeting Drosha Exon 4 was used for the generation of a Jurkat Drosha k.o. pool and editing efficiency of the sgRNA post-transfection and expansion of cells was determined to be 39% by Synthego suggesting that Drosha is required for fitness in Jurkat cells. Jurkat Drosha k.o. clonal cell lines #E11 and #F6 were isolated from the pool by limiting dilution and clonal expansion and cultured in suspension in RPMI1640 medium (Corning #10–040 CM), supplemented with 10% heat inactivated FBS (Sigma-Aldrich #14009C) and 1% penicillin/streptomycin (Corning #30-002-CI). Cell cultures were maintained at 37°C in a humidified 5% CO_2_ incubator (Nuaire, NU-5510).

### Production of HIV virus and viral infections

HIV virus was produced by transient transfection of HEK293T cells using Polyethyleneimine (PEI, 1 mg/ml, linear, MW 25000, Polysciences Inc.). Pseudotyping of GFP-HIV (HIV-1 NL4-3 ΔEnv EGFP Reporter Vector, a gift from Tom Hope, Northwestern University) by the vesicular stomatitis virus glycoprotein (VSV-G) was performed by cotransfection with pCMV-VSV-G vector. Briefly, transfection mixes were prepared using PEI (4 μl per 1 μg of DNA) with the GFP-HIV (6 μg) and VSV-G (4 μg) constructs in serum-free DMEM medium (Corning #10-013-CM) and added dropwise to HEK293T cells that were seeded onto 10-cm dishes a day prior at a density of 5-6 × 10^6^ cells in 10 ml of fresh DMEM with 10% heat inactivated FBS (Sigma-Aldrich #14009C). 18hrs after transfection, the medium was replaced with 10 ml of fresh DMEM with 10% heat inactivated FBS (Sigma-Aldrich #14009C). After 48 hrs the supernatant containing VSV-G pseudotyped HIV virus was collected and clarified by centrifugation at 450 x g for 5 min. and filtered through a sterile 0.45 μm-pore-size filter (MillexHV, Millipore). Viral supernatants were then used for infections or frozen in aliquots at −80°C.

For precipitation of the VSV-G pseudotyped HIV viral particles, 2.5 ml of sterile PEG-it solution (System Biosciences, #LV810A-1) was added to 10 ml of filtered viral supernatant and stored at 4°C for 48 hrs. Following incubation at 4ºC, the supernatant/PEG-it mixture was centrifuged at 1500 x g for 30 mins at 4°C. The supernatant was removed and the vector particles which appeared as a pellet were resuspended in 1 ml of PBS. 200 μl of the sample was provided to the viral pathogenesis core (Third Coast CFAR) for a p24 ELISA to quantitatively measure HIV-1 p24 protein. PEG-it concentrated virus was frozen in aliquots at −80°C.

To study effects of HIV on cells, growth/viability assays and flow cytometry analyses were performed following infection of cells with either VSV-G pseudotyped HIV viral supernatant (1-5% of total media volume) or PEG-it concentrated virus (in ng p24/ml). Additionally, effect of HIV on cell types was also assessed in the presence of the non-nucleoside reverse transcriptase inhibitor (NNRTI) Efavirenz (EFA) (Sigma-Aldrich #SML0536) or the integrase strand-transfer inhibitor Raltegravir (RAL) (Abcam #ab231360). Replication competent HIV NL4-3 Nef:IRES:GFP (NIH AIDS Reagent Program, cat. no. 11349) virus was produced and cells were infected as described (76). J-Lat cells were activated to release viral particles using PMA (0.1 μM).

### Cell growth and viability assessment

To monitor cell growth following HIV infection, adherent cells were seeded at a density of 3 × 10^3^ cells in 100 μl media/well in a 96-well plate and infected with either supernatant (2.5%) or concentrated VSV-G pseudotyped HIV (25, 50 or 175 p24ng/ml in a total media volume of 200 μl. Samples were assessed in quadruplicates. Cell growth was then monitored (-/+ HIV) at various time-points for at least 160 hours using the IncuCyte Zoom live-cell imaging system (Essen Bioscience, RRID:SCR_019874) with a 10X objective and data management software (version 2015A) to monitor cell proliferation. In each well, two phase-contrast images or GFP images were taken and the mean ± SEM percent confluence or green object integrated intensity was calculated using software processing definitions for the cell lines as recommended by the manufacturer.

To monitor cell viability following HIV infection, adherent or suspension cells were seeded at a density of 4-5 × 10^3^ cells in 100 μl media/well in a white-opaque 96-well plate and infected with either supernatant or concentrated VSV-G pseudotyped HIV in a total media volume of 200 μl. Samples were assessed in quadruplicates. A cell-viability assay involving the measurement of ATP within cells -/+ HIV was performed 72 hrs following HIV infection unless otherwise indicated. Briefly on the day of cell-viability measurement for adherent cells, media in each well was replaced with 70 μl of fresh media and 70 μl CellTitre-Glo reagent (Promega #G7570). For suspension cells, the plate was briefly centrifuged (1500 rpm for 30 seconds) to ensure all cells were settled at the bottom of the plate, followed by replacing 100 μl of culture media in each well with 100 μl CellTitre-Glo reagent (Promega #G7570). Plates were covered with aluminum foil and placed on an orbital shaker to induce cell-lysis for 5 min. followed by incubation at room temperature for 10 minutes before quantifying luminescence on a BioTek Cytation 5 cell-imaging multimode reader. The mean luminescence and standard deviation of the replicates for each condition was calculated in Microsoft Excel 2016 (Microsoft Corporation, USA) and represented as a percentage of the mean luminescence of the uninfected controls designated as 100% viability. Percent viability was then plotted against media control or HIV infection and represented in a Bar Graph format.

Jurkat derived cells were seeded at a density of 1 × 10^5^ in 200 μl media/well in a 96-well U-bottom plates and infected with replication competent HIV NL4-3 Nef:IRES:GFP at MOI 2, 5 and 10 in a media volume of 250 μl. Cell viability following HIV-infection was monitored at different time-points 72 hrs, 6 days and 8 days by transferring 50 μl of infected cells from the original culture plate to a fresh white-opaque 96-well plate containing 50 μl culture media. Each condition was assessed for cell viability in triplicate. 100 μl of CellTitre-Glo reagent (Promega #G7570) was then added to each well and luminescence was measured to quantify viable cells as described above.

J-Lat cells were seeded at a density of 5 × 10^3^ cells/well in a white-opaque 96-well plate and treated with either DMSO control or PMA (0.1 μM) in a total media volume of 200 μl. Samples were assessed in triplicates. Cell viability was assessed 48 and 96 hrs following PMA treatment by brief centrifugation (1500 rpm for 30 seconds) of the plate to ensure all cells are settled at the bottom of the plate, followed by replacing 100 μl of culture media in each well with 100 μl CellTitre-Glo reagent (Promega #G7570). Luminescence was measured to quantify viable cells as described above.

To monitor cell growth following transfection with short oligonucleotides, 50 μl Optimem (Thermo Fisher Scientific #31985070) mix containing 0.3 μl of RNAimax (optimized for HCT116 derived cells) and 1 or 10 nM miRNA or siRNAs was added per well of a 96-well plate. 3000 cells in 200 μl antibiotic free medium were added to each well containing the transfection mix. Each transfection condition including test miRNA/siRNA, negative control and RNAiMax control was performed in quadruplicates and growth was monitored as described above in a IncuCyte Zoom live-cell imaging system (Essen Bioscience, RRID:SCR_019874).

The following miRNAs and negative controls were used: miR-TAR-3p (mirVana™ miRNA Mimic, mature sequence: UCUCUGGCUAACUAGGGAACCCA, Ambion Cat. No# 4464066), S1_TAR-3p (mirVana™ miRNA Mimic, mature sequence: UGGCUAACUAGGGAACCCACUG, Ambion Cat. No# 4464068), S2_TAR-3p (mirVana™ miRNA Mimic, mature sequence: CUCUGGCUAACUAGGGAACCCACU, Ambion Cat. No# 4464068) and NC1 (miRNA precursor negative controls #1, Ambion, Cat. No# AM17110). The following siRNA oligonucleotides were designed as described in (77) and obtained from Integrated DNA Technology and annealed as per the manufacturer’s instructions:

siNT1modified sense: mUmGrGrUrUrUrArCrUrArCrArCrGrArCrUrArUTT;

siNT1modified antisense: rArUrArGrUrCrGrUrGrUrArGrUrArArArCrCrAAA

siTAR-3p (2-7) sense: mUmGrGrUrUrUrArCrArUrGrUrCrCrUrGrArGrATT

siTAR-3p (2-7) antisense: rUrCrUrCrUrGrGrArCrArUrGrUrArArArCrCrAAA

siTAR-3p (3-8) sense: mUmGrGrUrUrUrArCrArUrGrUrGrCrCrArGrArATT

siTAR-3p (3-8) antisense: rUrUrCrUrGrGrCrArCrArUrGrUrArArArCrCrAAA

si-S1_TAR-3p (2-7) sense: mUmGrGrUrUrUrArCrArUrGrUrUrUrArGrCrCrATT

si-S1_TAR-3p (2-7) antisense: rUrGrGrCrUrArArArCrArUrGrUrArArArCrCrAAA

si-S1_TAR-3p (3-8) sense: mUmGrGrUrUrUrArCrArUrGrUrGrUrUrArGrCrATT

si-S1_TAR-3p (3-8) antisense: rUrGrCrUrArArCrArCrArUrGrUrArArArCrCrAAA

### Cell-death assay

1 × 10^6^ cells were infected with VSV-G pseudotyped HIV R9 ΔEnv iGFP (175 ng p24/ml) in the absence or presence of antivirals (10 μM EFA or 10 μM RAL). 72 h following infection, cells were harvested and stained using LIVE/DEAD fixable Blue Dead Cell Stain (Invitrogen #L23105), according to manufacturer’s instructions. HIV infected cells and cell viability were both analyzed by Flow cytometry (BD LSRFortessa) and quantified using FlowJo™ v10.

### Cellular surface staining and flow cytometry analysis

To study the expression of the surface receptors LDL-R, CD4 and CXCR4, cellular surface staining was performed. Briefly 6 × 10^5^ cells were pelleted and washed in 1x PBS and resuspended in 100 μl of FACS buffer (PBS+ 10% FBS + 0.1% sodium azide) in a microfuge tube. Duplicates for each condition were set up, where 20 μl of isotype control antibody or surface receptor specific primary labelled antibody was added to the 100 μl cell suspension in FACS buffer and then incubated on ice in the dark for 30 mins. Following incubation, the cells were washed with 1 ml FACS buffer 3 times by centrifugation at 1500 rpm for 5 min. and then resuspended in 500μl of ice cold FACS buffer. The cell suspension was then transferred to FACS tubes and placed on ice in the dark and then immediately taken for analysis by flow cytometry (BD FACSymphony-A5). Percent surface receptor positive cells was quantified by FlowJo™ v10. The following antibodies were used: PE Mouse Anti-Human LDLR (BD Biosciences, Cat No. 565653) and PE Mouse IgG2b, k Isotype control (BD Biosciences, Cat No. 555058); PE-Cy5 Mouse Anti-Human CD184 (BD Biosciences, Cat No. 555975) and PE-Cy5 Mouse IgG2a, κ Isotype control (BD Biosciences, Cat No. 555575); APC Mouse Anti-Human CD4 (BD Biosciences, Cat No. 561840) and APC Mouse IgG1, κ Isotype control (BD Biosciences, Cat No. 555571).

### Western blot analysis

Protein lysates were prepared by lysing cells using Killer RIPA lysis buffer (150 mM NaCl, 10 mM Tris HCl pH 7.2, 1% sodium dodecyl sulfate (SDS), 1% TritonX-100, 1% Deoxycholate, 5 mM EDTA) supplemented with PMSF (1 mM) and Protease Inhibitor Cocktail (Roche #11836170001) followed by sonication of samples (25 % Amplitude for 10 seconds). DC Protein Assay kit (Bio-Rad, Hercules, CA) was used to quantify protein concentration in each sample. Equal amounts of protein per sample (25-30μg) were resolved on a 10% SDS-PAGE gel and transferred onto nitrocellulose membranes (Amersham Protran 0.45 μm NC, GE) at 90V for 90 min. Membranes were incubated in blocking buffer (5% dry milk in 0.1% TBS/Tween-20) for 1 hour at room temperature. Appropriate dilution of primary antibodies was prepared in blocking buffer and applied to membranes and incubated overnight at 4°C. Membranes were washed three times in 0.1% TBS/Tween-20 for 10 min. each. Secondary antibodies were diluted in blocking buffer and applied to membranes for 1 hr at room temperature followed by three additional washing steps with 0.1% TBS/Tween-20 for 10 min. each. The protein-antibody complexes were detected using the SuperSignal™ West Dura Extended Duration Substrate (Thermo Fisher Scientific #34076) and visualized using the chemiluminescence imager G:BOX Chemi XT4 (Syngene). The following primary antibodies used were diluted in blocking buffer: anti-Argonaute-2 (1:1000) (Abcam #32381), anti-Argonaute-1 (1:1000) (Cell Signaling #5053), anti-Argonaute-3 (1:1000) (Cell Signaling #5054), anti-Argonaute-4 (1:1000) (Cell Signaling #6913), anti-Dicer (1:1000) (Cell Signaling #D38E7), anti-Drosha (1:1000) (Cell Signaling #D28B1), anti-Exportin 5 (1:1000) (Cell Signaling #12565), anti-β-actin (1:5000) (Santacruz #Sc-47778). The secondary antibody Goat anti-rabbit IgG-HRP (1:5000) (Southern Biotech #4030–05) was diluted in blocking buffer. Densitometry analysis of the blots were carried out using GeneTools (Syngene) image analysis software and protein bands were normalized to the loading control-β-actin. All uncropped Western blots are shown in **Fig. S8**.

### Ago pull-down and small RNA-sequencing (Ago-RP-Seq)

To determine the RISC content of cells infected with HIV, an Ago pull-down experiment followed by high-throughput small RNA sequencing was performed. All experiments involving HIV infection described above for 96-well plates were scaled up for 150 mm dishes. Briefly, Jurkat derived cells were infected with 50 ng p24/ml of PEG-it concentrated VSV-G pseudotyped HIV and HCT116 derived cells were infected with either 175 ng p24/ml of concentrated VSV-G pseudotyped HIV or 2.5% of viral supernatant and harvested for pull-down 28 hrs following infection. Jurkat derived cells infected with replication competent HIV NL4-3 Nef:IRES:GFP at MOI 10 were harvested 12 days following infection and J-Lat cells were harvested 36 hrs following induction with 0.1 μM of PMA.

Harvested cells were washed with 1x PBS, and 10^6^ cells were pelleted and frozen at -80°C until ready to be lysed with 1 ml NP40 lysis buffer [50 mmol/L Tris pH 7.5, 150 mmol/L NaCl, 5 mmol/L EDTA, 0.5% NP-40, 10% (v/v) glycerol, 1 mmol/L NaF; supplemented with 1:200 EDTA-free protease inhibitors (Millipore #539134) and 1:1000 RNaisin Plus (Promega #N2615) before use]. Lysates were incubated on ice for 15 min. and vortexed briefly followed by centrifugation at 20,000 x g for 20 mins at 4°C. Lysates were transferred to siliconized microcentrifuge tubes (DNA LoBind, Eppendorf #022431021), and 500 μg of Flag-GST-T6B peptide with high-affinity binding to Ago1-4 (78) was added coupled with 80 μl of anti-Flag M2 Magnetic beads (Sigma #M8823) and incubated for 3 hrs on a rotor at 4°C. The beads were then washed three times with NP40 lysis buffer and during the final wash, 10% of the beads were removed and incubated at 95°C for 5 minutes with 4x Laemmli sample buffer added. Ago pull-down efficiency was assessed by running these samples on a 10% SDS-PAGE gel, transferring onto a nitrocellulose membrane and immunoblotting against Ago2 (Abcam #32381). 500 μl of TRIzol reagent was added to the remaining beads and a phenol: chloroform RNA extraction was performed according to manufacturer’s instructions. The RNA pellet was dissolved in 20 μl RNAse free water. 10 μl of the RNA sample was dephosphorylated with 0.5 U/mL of CIP alkaline phosphatase at 37°C for 15 minutes and then radiolabeled with 0.5 mCi [γ-^32^P] ATP and 1 U/ml T4 PNK kinase for 20 min. at 37°C. RNA bound to Ago 1-4 was visualized by 15% Urea-PAGE. The remaining RNA sample was used for small RNA library preparation using Illumina primers (RRID:SCR_010233) as described previously (79). Briefly, RNA was first ligated with 3’-adenylated adapters and separated by 15% denaturing Urea-PAGE. The RNA corresponding to insert size of 19–35 nt was eluted from the gel using P^32^ radiolabeled size markers, ethanol precipitated and then ligated with the 5’ adapter. The RNA samples were then separated by 12% Urea-PAGE followed by gel extraction and subsequent reverse transcription using Superscript III reverse transcriptase (Invitrogen #18080–044). The cDNA was PCR amplified and sequenced on Illumina Hi-Seq 4000. The sequences used for the RNA size marker, 3’- and 5’-adaptors, RT primers and PCR primers were used as described previously (80).

### 6mer seed toxicity analysis of R-sRNAs

The SPOROS pipeline was used to analyze the abundance of RISC bound sRNAs according to their predicted 6mer seed toxicity and generate seed toxicity graphs and seed toxicity plots (81). In brief, the RISC bound small RNA Seq data were trimmed to remove adaptor sequences and reads that are longer than 25nt or shorter than 18nt and complied into a counts table. Reads with fewer counts than the number of samples were removed, and the remaining reads were BLASTed against a curated list of all mature human miRNAs or other short RNAs (RNAworld). Reads that matched any artificial sequences in the RNA datasets were also removed and the subsequent raw read count table (RawCounts) was either normalized to 1 million reads to generate a norm count table (normCounts) or normalized and used for differential expression analysis (differential). Each table was then annotated with 6mer seed, predicted 6mer seed viability (as determined from https://www.6merdb.org/), miRNA and RNAworld matches. At this step, the percent miRNA/short RNA content in the RISC was determined from the annotated normCount table. Seed toxicity graphs were then generated by collapsing the table according to 6mer seed and RNA type and aggregating all the rows according to the predicted 6mer seed toxicity in 1% sized bins and then adding up all the seeds that had a specific toxicity with each 1% bin. Line graphs with smoothing enabled were then generated in Microsoft Excel, with 6mer seed viability on the x-axis and normalized read count on the y-axis. The data file with the collapsed data was used to identify the most abundant miRNAs/short RNAs that make up each peak of the seed tox graph and was labelled if >5000 reads. Where a peak consisted of more than one abundant miRNA/short RNA i.e., >5000 reads, RNAs were listed in order of abundance. To determine average seed viability of all reads in the data set, the counts for all individual reads were divided by a factor that reduces the number of the most abundant reads to <1000 to ease computational complexity of the downstream analyses. The columns containing the 6mer seed viability were then expanded according to the normalized read counts and the resultant data column containing the frequency distribution of the average 6mer seed viability for various groups was used to generate a basic boxplot (Seed tox plot) using StatPlus (v 7.7)). Differential analysis between two groups was performed by taking significantly (p-value <0.05) up or downregulated reads and calculating the delta read counts for each row between groups (perturbed sample-control sample). Associated 6mer seed viability were then multiplied based on the delta read counts to produce the seed toxicity graphs and seed toxicity plots as described above.

### Identification of HIV derived reads in the RISC of HIV-infected cells

Briefly, the reads from each sample were compiled as a BLAST database, and blastn was utilized to query the HIV-1 whole genome sequence (NCBI Reference Sequence: NC_001802.1) against reads from cells infected with HIV. Reads were considered matches if they had an e-value < 0.05% and 100% identity across the entire length of the read. The filtered BLAST hits were converted to a bed formatted file, describing the locations of reads relative to the HIV-1 whole genome and was used for subsequent analyses. Sample replicates were combined and stack plots depicting the locations where RISC bound HIV sRNAs map along the HIV-1 whole genome were generated using R package Sushi. HIV sRNA reads containing the core region of the HIV-miR-TAR3p sequence-UGGCUAACUAGGGAAC were highlighted in red.

### Transfection of cells with miR-TAR-3p and large RNA-seq analysis

HCT116 wt cells were seeded at a density of 2.5 x10^5^ cells/well in 6-well plates. The following day, cells were forward transfected with miR-TAR-3p (mirVana™ miRNA Mimic, mature sequence: UCUCUGGCUAACUAGGGAACCCA, Ambion Cat. No# 4464066) or NC1 (miRNA precursor negative controls #1, Ambion, Cat. No# AM17110) in duplicates with 500 μl Optimem mix containing 3 μl RNAimax and miRNA mimic at 10 nM was added to 2 ml of antibiotic free medium. Cells were harvested for RNA 24 hrs following transfection using Qiazol. An on-column DNAse digestion step was included using the RNAse-free DNAse set (Qiagen #79254) and total RNA was isolated using the miRNeasy^®^ Mini Kit (Qiagen # 217004) according to manufacturer’s instructions. The quality of RNA was determined using an Agilent Bioanalyzer. RNA library preparation and subsequent sequencing on NovaSeq SP PE150 was performed by the NU-Seq core at Northwestern University (Chicago). Paired-end RNA-seq libraries were prepared using Illumina TruSeq Total RNA library prep kit with ribozero rRNA depletion step included. Reads were trimmed with Trimmomatic v0.33 (TRAILING:30 MINLEN:20) (82) and then aligned to the hg38 human genome assembly with Tophat v 2.1 (83) or STAR (84). Exonic reads were assigned to genes using the Ensembl 78 version of the hg38 transcriptome and htseq v 0.6.1 (85). Differential expression analysis was carried out using the edgeR package (86) to fit a negative binomial generalized log-linear model to the read counts for each gene. Sylamer analysis was performed as recently described (38). 3’ UTRs or ORFs were used from Ensembl, version 76.

### miRNA quantification

Briefly, 25 ng total RNA was used to make cDNA using the High-Capacity cDNA Reverse Transcription Kit (Applied Biosystems #4368814) and miRNA specific primers. The cDNA was diluted 1:5 and a qPCR reaction mixture composed of the diluted cDNA, target specific TaqMan microRNA assay (Thermo Scientific #4427975), and the Taqman Universal PCR Master Mix (Applied Biosystems #4324018) was prepared. Ct values were determined as described previously using the Applied Biosystems 7500 Real Time PCR system. The ΔΔCt values between the miRNA of interest and the control were calculated to determine relative abundance of the miRNA. Each sample was run in triplicate and z30 (Thermo Scientific Assay ID #001092) was used as an endogenous loading control. The following TaqMan microRNA assays (Thermo Scientific #4427975) were used: miR-21 (Assay ID: 000397), miR-200b (Assay ID: 002251), and miR-320 (Assay ID: 002277).

### Ago4 knockdown assessment

2.5 × 10^5^ cells of HCT116 and Ago123 TKO cells were reversed transfected with 40 nM siNT (Dharmacon ON-TARGETplus non-targeting pool #D-001810-10-05) and siAgo4 (Dharmacon ON-TARGETplus Human AGO4 siRNA # L-004641-01-0005) in a 6-well plate with 3 μl RNAimax. A mock condition with RNAimax was also set up for each cell type. Cells were harvested for RNA 48 hrs following transfection using the miRNeasy^®^ Mini Kit (Qiagen # 217004) according to manufacturer’s instructions and 200 ng RNA was used to make cDNA using the High-Capacity cDNA Reverse Transcription Kit (Applied Biosystems #4368814). The cDNA was diluted 1:5 and a qPCR reaction mixture composed of the diluted cDNA, Ago4 (Thermo Scientific Assay ID: Hs01059739_m1) or GAPDH (Thermo Scientific Assay ID: Hs00266705_g1) specific TaqMan primer-probe, and the Taqman Universal PCR Master Mix (Applied Biosystems #4324018) was prepared. Ct values were determined as described previously using the Applied Biosystems 7500 Real Time PCR system. Each sample was run in triplicate and the ΔΔCt values between the gene of interest and control were calculated to determine relative abundance of the gene. Samples were normalized to GAPDH.

### Statistical analysis

Statistical analysis was performed using Student’s T-test. P values for statistical significance between the groups of deregulated sRNAs (in seed toxicity plots) were determined using a Kruskal–Wallis median rank test. Two-way ANOVA was performed using the Stata1C software for evaluating GFP positivity over increasing HIV concentrations, choosing one component for the treatment condition as primary interest and a second component for the drug concentration as a categorical variable.

## Acknowledgements

We are grateful to David Corey and Joshua Mendell for providing the Ago1 k.o. and Ago123 k.o., and Ago2 k.o. HCT116 cells, respectively. This work was supported by NIH grant R21 AI150910 to M.E.P.

