## Supplementary figures for "Contribution of 6mer seed toxicity to HIV-1 induced cytopathicity"

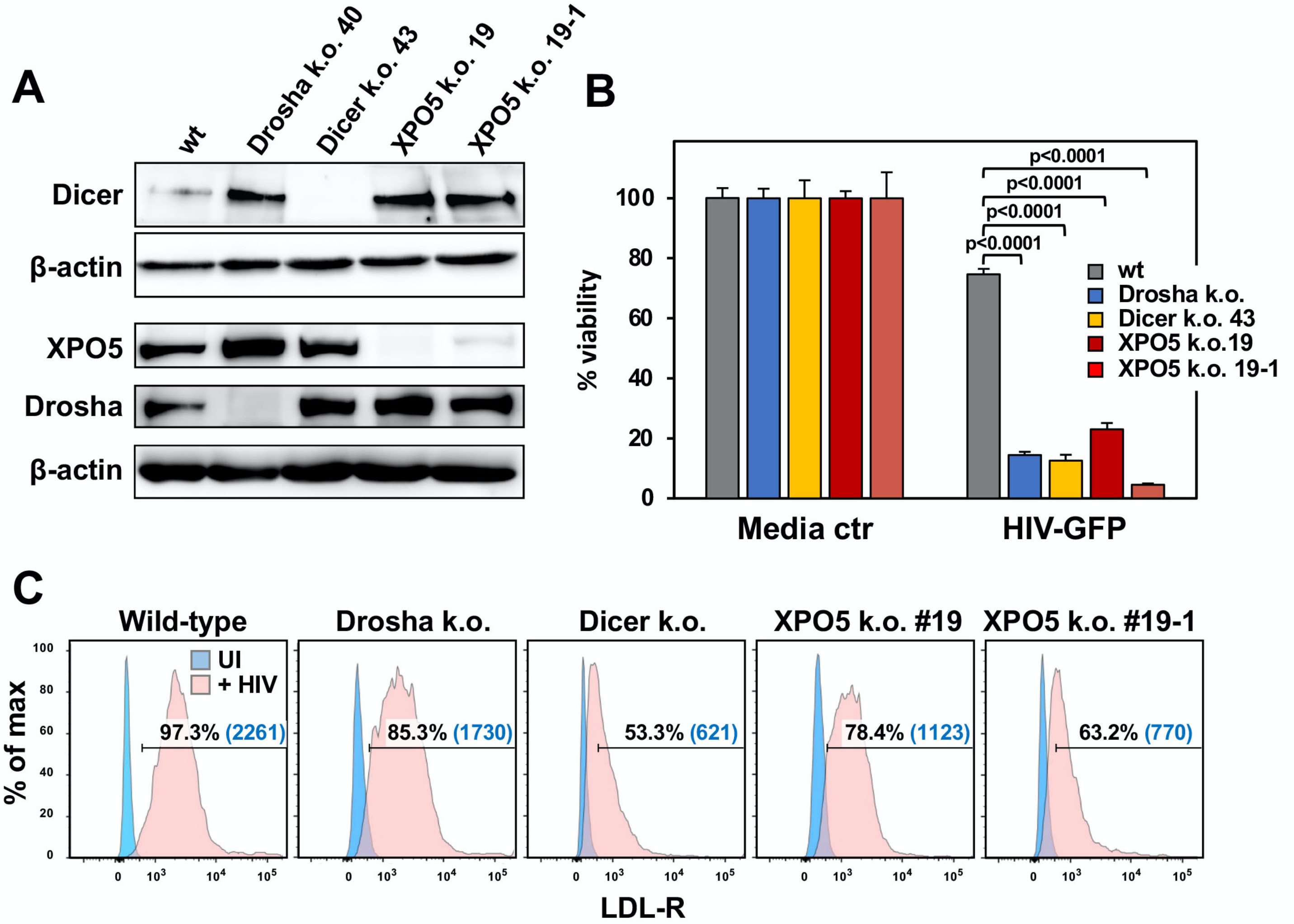


**Figure S1: Characterization of different mutant HCT116 cells.**
(**A**) Western blot analysis of wt HCT116 cells, Dicer k.o., Drosha k.o., or two XPO5 k.o. HCT116 clones.
(**B**) Viability of the HCT116 wt and mutant cells determined 72 hours after infection with 2.5% of VSV-G pseudotyped HIV supernatant. Student's t-test p-values are displayed.
(**C**) Surface staining of LDL-R in all cell lines shown in A. Percent positivity and median fluorescence intensity (in brackets) is shown.

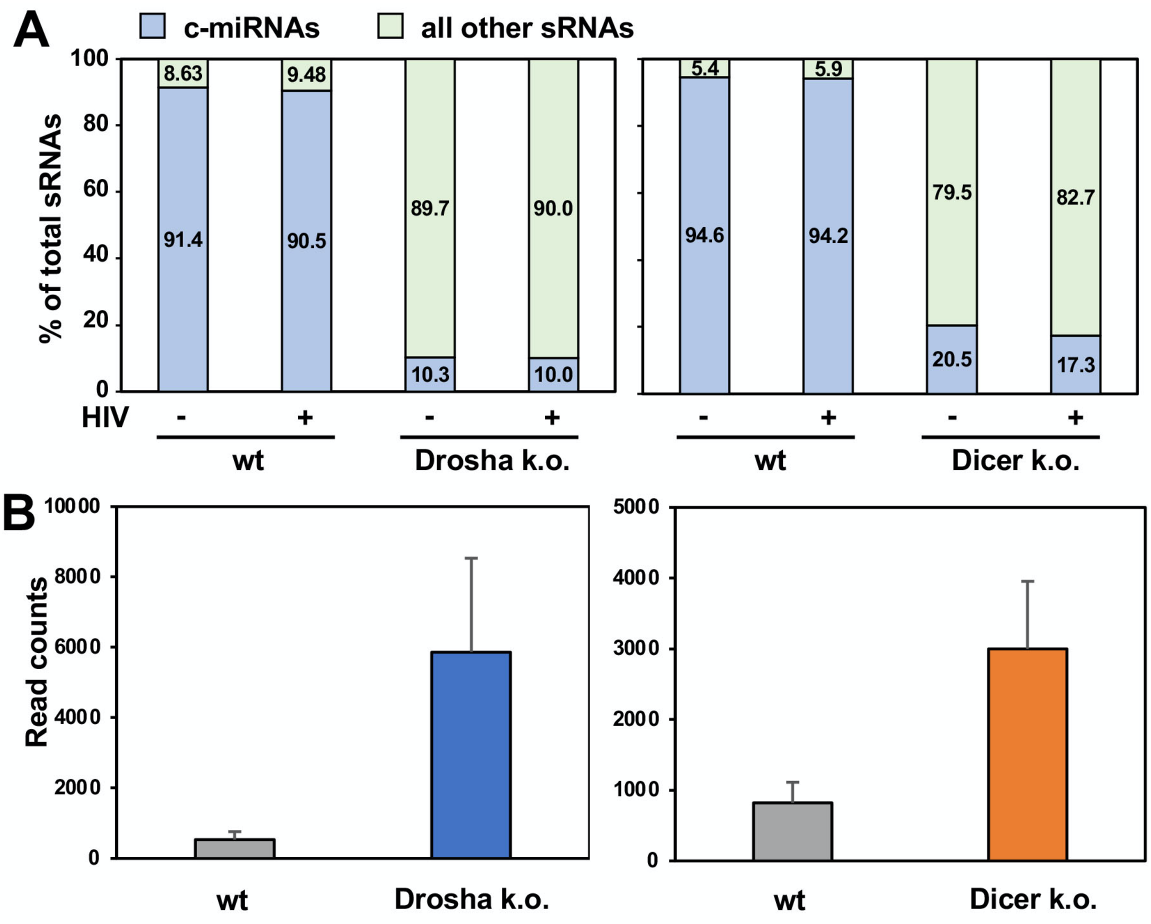


**Figure S2: Overall effects of HIV infection on miRNA content in HCT116 wt, Drosha, or Dicer k.o. cells**
(**A**) Percent c-miRNAs bound to RISC of wt and Drosha k.o. cells (left) or wt and Dicer k.o. (right) cells uninfected (-) or infected (+) with VSV-G pseudotyped HIV.
(**B**) RawCounts of viral R-sRNAs in cells as described in A.

**A**

**5' arm**

```
C**TCTGGT**TAGACCAGATCTGAGCCT
 T**CTGGTT**AGACCAGATCTGAGCCT
  C**TGGTTA**GACCAGATCTGAGCCT
  C**TGGTTA**GACCAGATCTGAGCC
   T**GGTTAG**ACCAGATCTGAGCC
```

**3' arm**

```
T**CTCTGG**CTAACTAGGGAACCC
T**CTCTGG**CTAACTAGGGAACC
T**CTCTGG**CTAACTAGGGAAC
  T**CTGGCT**AACTAGGGAACCC
    T**GGCTAA**CTAGGGAACCCACT
```

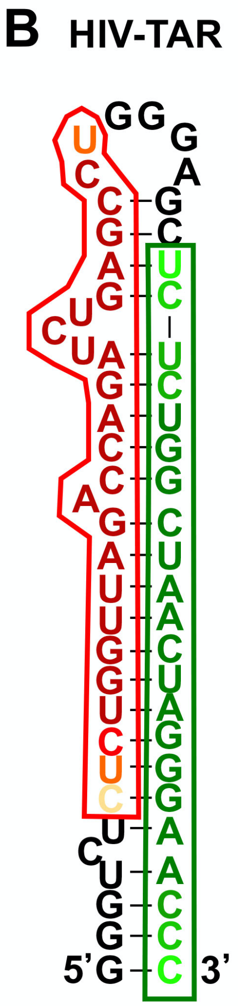


**Figure S3: HIV-TAR derived reads show potential for 6mer seed toxicity.**
(**A**) Different reads extracted from in HIV infected cells (1). The 6mer seeds of all HIV-TAR derived reads are highlighted in bold.
(**B**) Location of the reads in A in the HIV-TAR loop. Darker colors indicate higher overlap between the different reads on each arm.

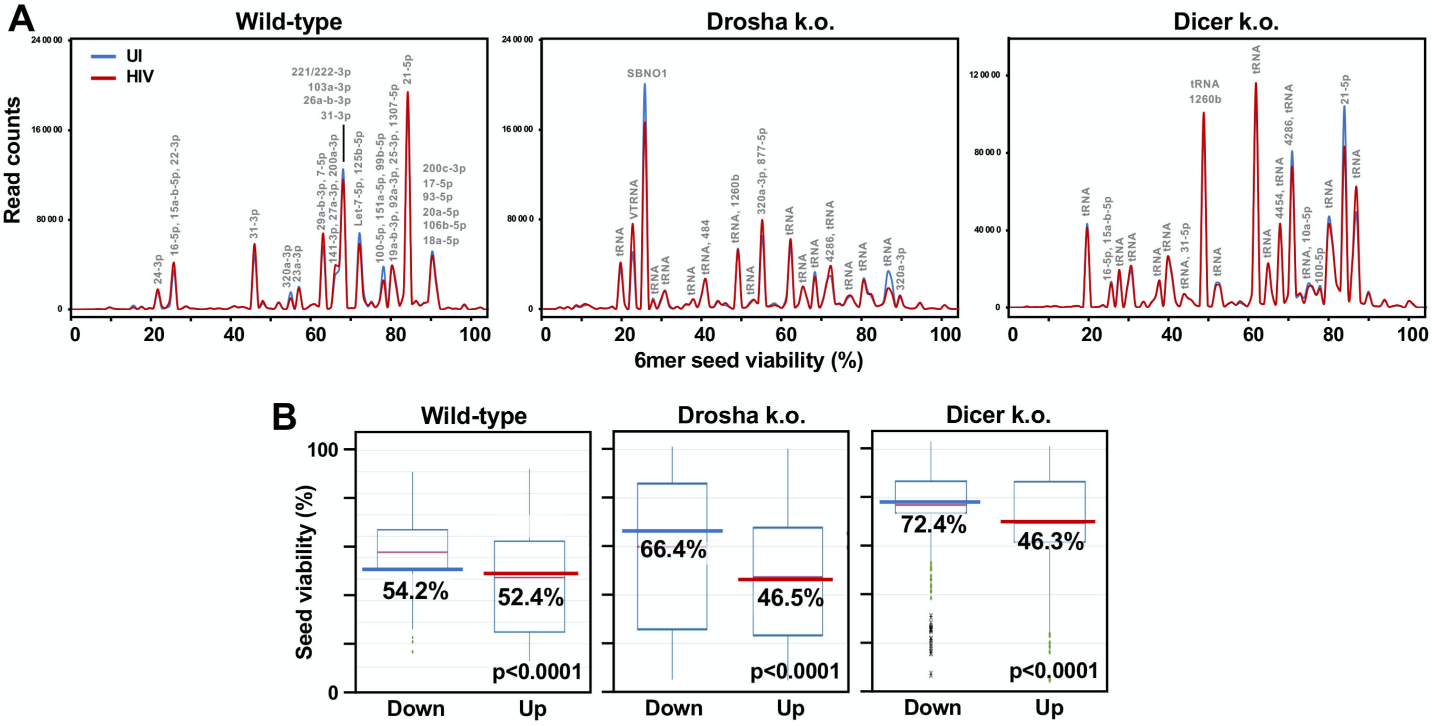


**Figure S4: Side-by-side comparison of seed viabilities of R-sRNAs in HIV infected HCT116 wt cells or Drosha or Dicer k.o. clones.**

(**A**) Seed toxicity graphs of all R-sRNAs in wt, Drosha, and Dicer k.o. cells uninfected or 28 hours after infection with VSV-G pseudotyped HIV. miRNAs that contribute to peaks with >5,000 reads are labeled. For each labeled peak, miRNAs are listed in the order of abundance.

(**B**) Seed toxicity box plots showing the median seed viability (%) of R-sRNAs significantly up or downregulated (>1.5 x, p< 0.05) in wt, Drosha k.o. or Dicer k.o. cells after HIV infection. The median is highlighted. P values of deregulated seed viabilities were determined using a Kruskal–Wallis median rank test.

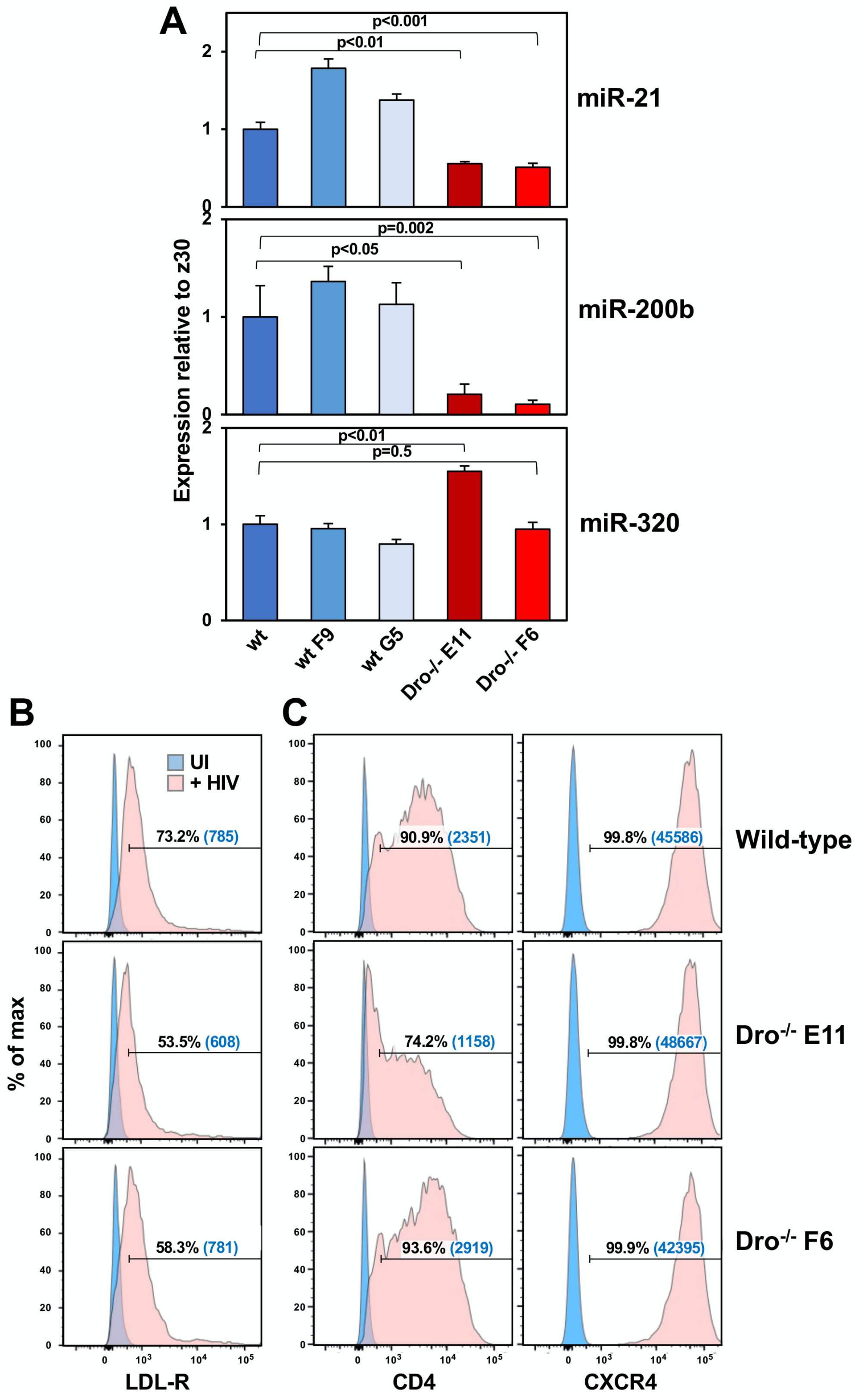


**Figure S5: Characterization of Drosha k.o. Jurkat clones.**
(**A**) Real-time PCR analysis of miRNAs in wt Jurkat cells and single cell (two wt and two Drosha k.o.) clones. miRNA expression was calculated relative to the expression of the z30 RNA. Student's t-test p-values are shown. (**B, C**) Surface staining of LDL-R (B), CD4 and CXCR4 (C) in wt Jurkat cells and the two Drosha k.o. clones. Percent positivity and median fluorescence intensity (in brackets) are displayed.

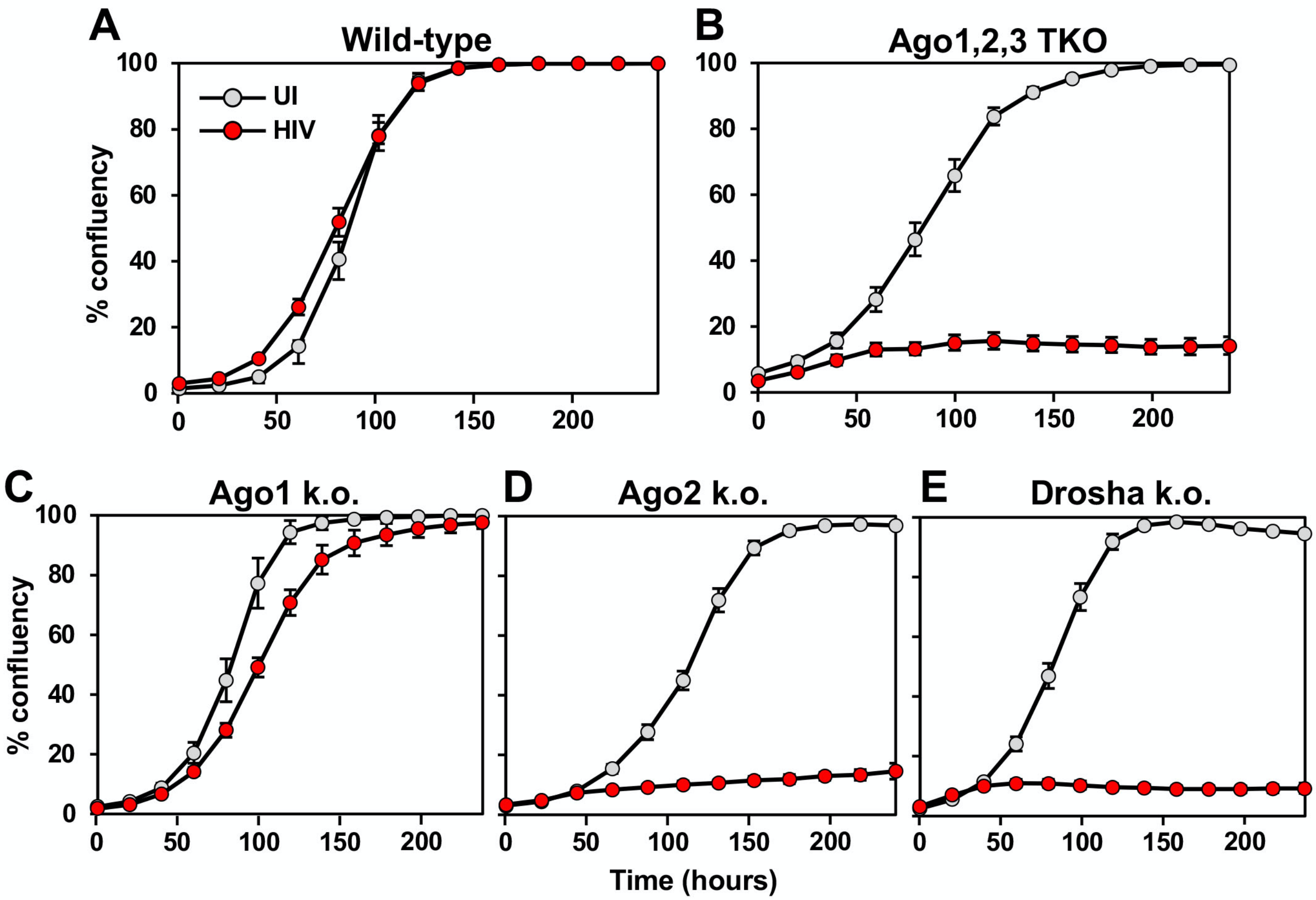


**Figure S6: Deletion of Ago2 results in much higher cell death induced by HIV.**
(**A-G**) Confluency over time of HCT116 wt cells or k.o. clones uninfected or infected with 1% (A-C, E) or 2.5% of VSV-G pseudotyped HIV supernatant (D). VSV-G pseudotyped HIV supernatant batch used in D had lower activity.

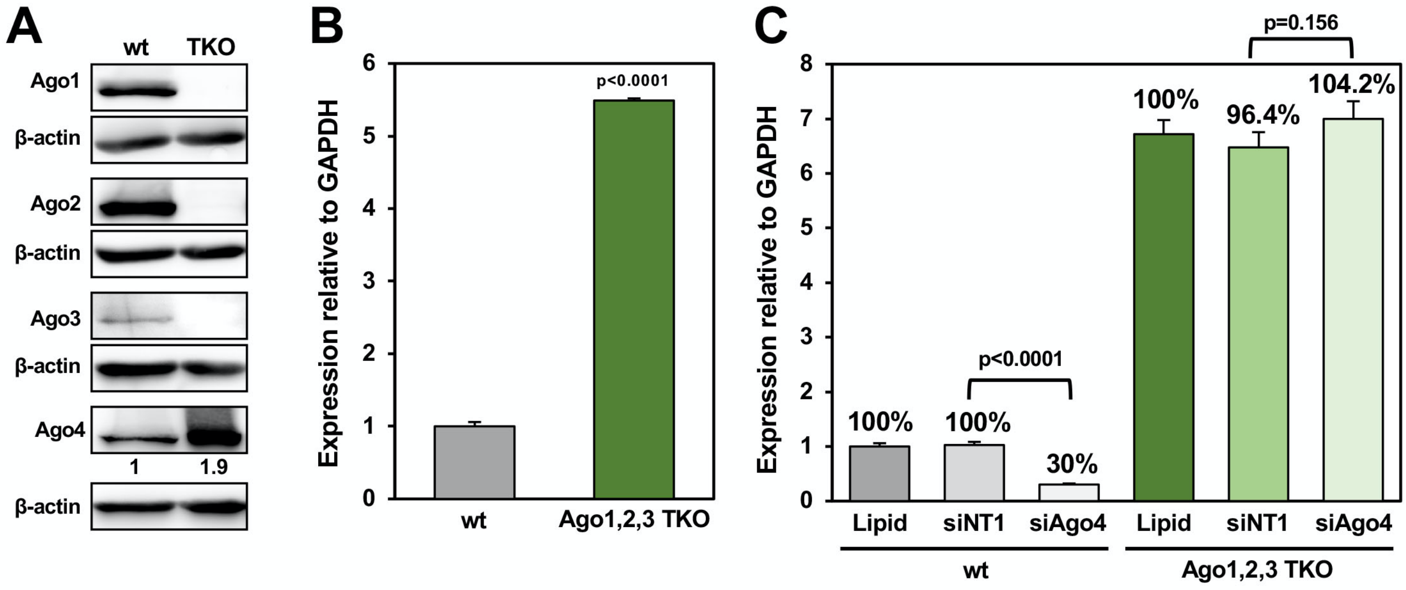


**Figure S7: A possible contribution of Ago4 to HIV induced toxicity in Ago1,2,3 TKO cells.**
(**A**) Western blot analysis of HCT116 wt and Ago1,2,3 TKO cells. Ago4 expression was quantified by densitometry relative to Actin.
(**B**) Real-time PCR analysis of Ago4 mRNA relative to GAPDH.
(**C**) Real-time PCR analysis of the Ago4 mRNA relative to GAPDH in HCT116 wt or Ago1,2,3 TKO cells 48 hours after transfection with a control SmartPool (siNT) or a SmartPool specific for Ago4.
Student's t-test p-values are displayed.

**Figure 4A**

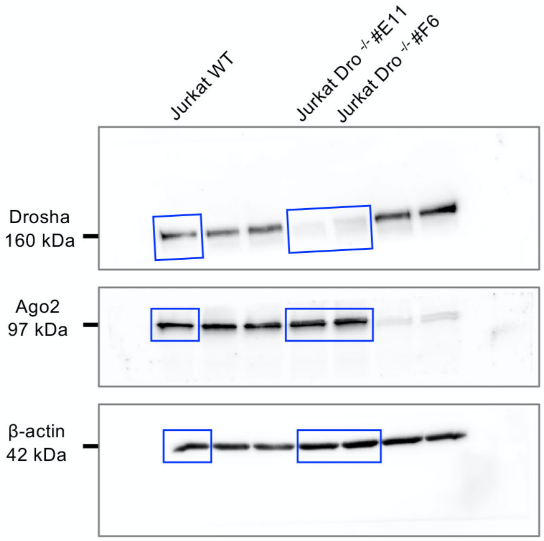


**Figure S1A**

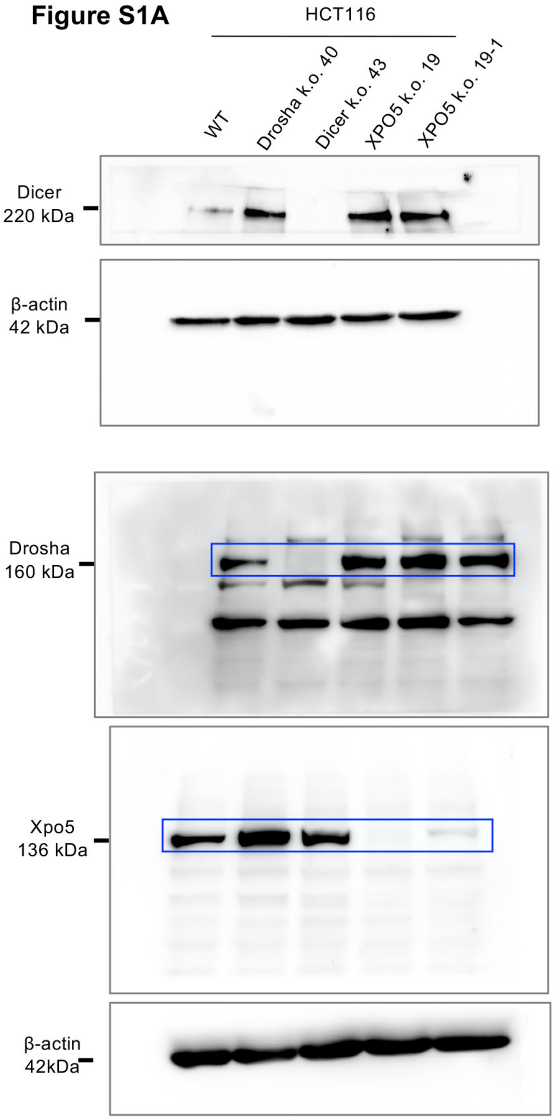


**Figure S7A**

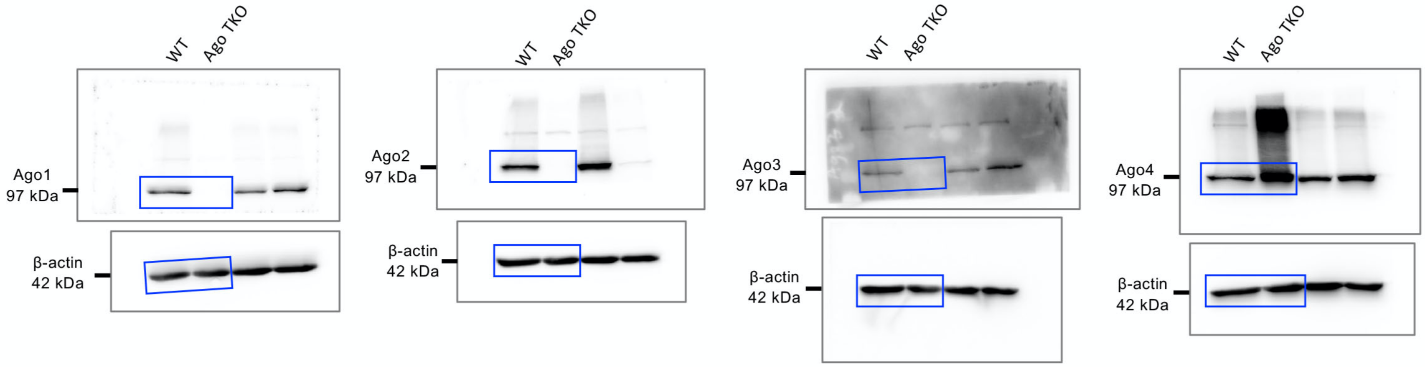


**Figure S8:** All uncropped Western blot images. Molecular weights for each blot are given in kDa. Regions used in the figures are boxed.
